# Viral transcriptional regulators extensively rewire host pathways through diverse mechanisms

**DOI:** 10.64898/2025.12.01.691387

**Authors:** Jaice T. Rottenberg, Xing Liu, Anna Berenson, Luis F. Soto-Ugaldi, Mohamed Y. ElSadec, Clarissa Santoso, James E. Corban, Phillip J. Dexheimer, Berkay Engin, Ryan Lane, Sakshi Shah, Kerstin Spirohn-Fitzgerald, Shubham Khetan, Cheng-Che Lee, George Munoz-Esquivel, Zhaorong Li, Lucia Martinez-Cuesta, Yunwei Lu, Philipp Trollmann, Tong Hao, S. Stephen Yi, Nidhi Sahni, Martha L. Bulyk, Michael Calderwood, Matthew T. Weirauch, Marc Vidal, Srivatsan Raman, Juan I. Fuxman Bass

## Abstract

Viral transcriptional regulators (vTRs) reprogram host gene regulatory networks to promote replication, persistence, and immune evasion. Despite the identification of hundreds of vTRs in human viruses, how they rewire host pathways remains unclear. Here, we systematically profiled 95 vTRs from diverse human viruses across multiple functional assays. vTRs perturb immune, cell proliferation/death, and signaling pathways through various mechanisms; some bind DNA directly, others cooperate or antagonize human transcription factors (hTFs), and some remodel chromatin. vTRs can act as activators or repressors and recruit similar but not identical repertoires of proteins as hTFs. These findings reveal vTRs as versatile transcriptional modulators that converge on conserved host “pressure points” while diversifying across pathways to promote viral replication and persistence. Notably, many vTR dysregulate genes within autoimmune, neurological, and cardiovascular risk loci, revealing mechanistic links to disease. Together, we provide a comprehensive resource for understanding and targeting viral control of human transcription.

## Introduction

Viruses are a leading cause of human disease, through direct effects of acute infection and by increasing the risk of chronic conditions, such as multiple sclerosis, dementia, or cancer.^1–5^ A key mechanism underlying viral pathogenicity is the ability to reprogram host gene expression to promote cell states suitable for viral replication or persistence.^6^ In particular, viruses often suppress immune response genes to evade detection and clearance, as well as modulate pathways controlling cell survival and proliferation to prevent elimination of infected cells and enhance viral spread.^7–10^ This viral manipulation of host transcriptional programs likely plays a central role in disease development and progression.

Viral proteins can modulate host gene expression either directly–by recruitment to cis-regulatory elements (CREs) in the host genome–or indirectly, by altering signaling pathways that control downstream genes.^6^ Direct modulation typically involves proteins classified as viral transcription regulators (vTRs), which, beyond controlling viral gene expression, can influence host transcription by binding DNA independently, co-binding with human transcription factors (hTFs), or being recruited to DNA via interactions with chromatin-associated proteins such as hTFs or cofactors (**Figure 1A**). For instance, the Epstein-Barr virus (EBV) Zta binds TPA-responsive elements to regulate immune genes,^11,12^ while HBZ from human T-lymphotropic virus 1 (HTLV-1) dimerizes with hTFs JUNB and JUN to modulate their DNA binding activity.^13^ In contrast, ChIP-seq studies have shown that vTRs such as EBV EBNA2 are indirectly recruited to human CREs through interactions with hTFs.^6^

**Figure 1:**
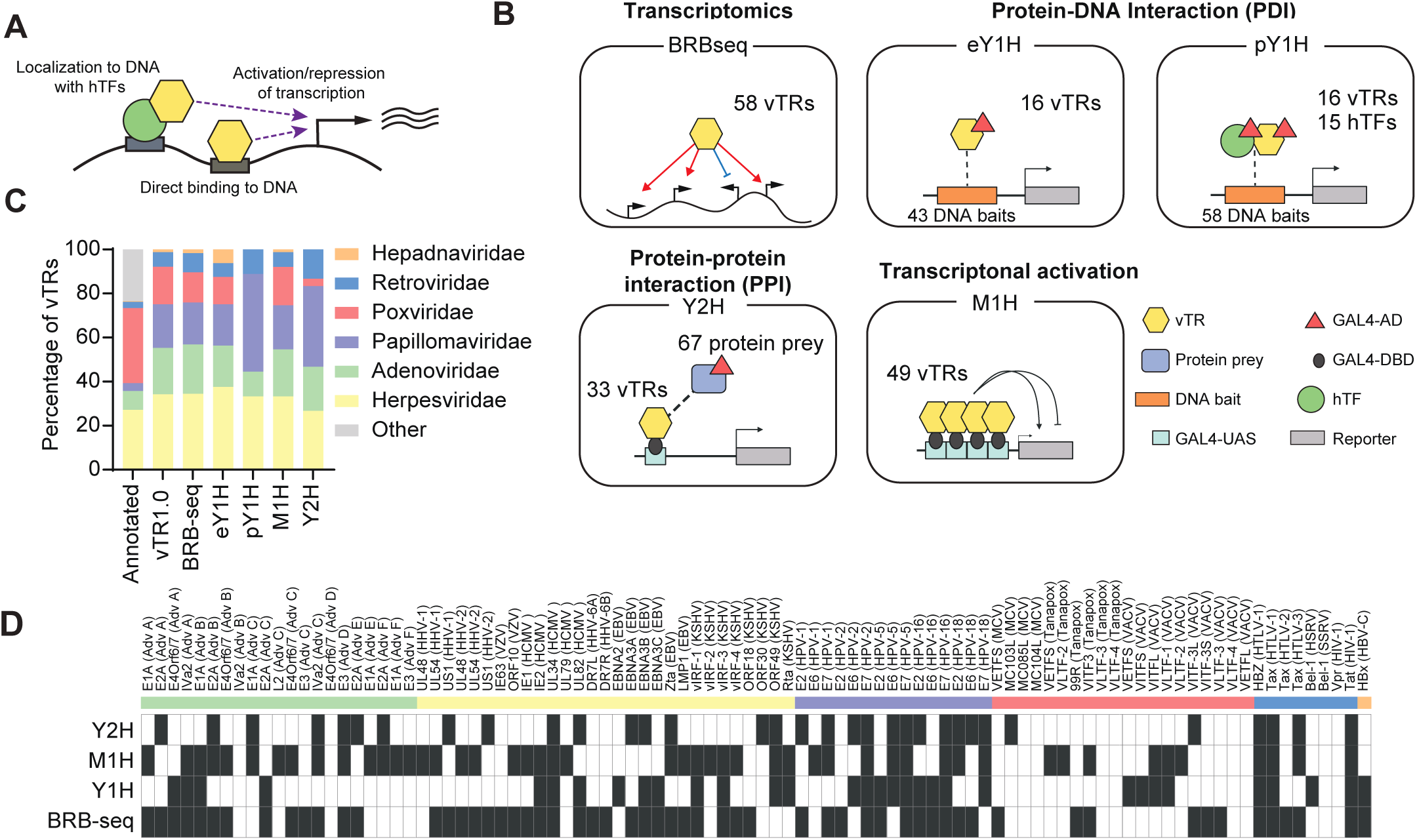
vTR1.0: a resource to study the viral transcriptional regulator functional landscape. (A) Schematic overview of vTR modulation of human transcription. (B) Overview of multimodal interrogation of vTR function. eY1H, enhanced yeast one-hybrid; pY1H, paired yeast one-hybrid; Y2H, yeast two-hybrid; M1H, mammalian one-hybrid. Gal4-AD, Gal4 activation domain; Gal4-DBD, Gal4 DNA-binding domain; Gal4-UAS, Gal4 upstream activation sequence; vTR, viral transcriptional regulator; hTF, human transcription factor. (C) Percentage of vTRs from different viral families represented across all annotated vTRs, the vTR1.0 resource, and tested in each experimental modality. (D) Data acquired for each vTR across experimental modalities. Columns correspond to vTRs (virus of origin in parenthesis). Rows correspond to each experimental modality. Black cells indicate data generate for a given vTR in each modality (based on parameters and thresholds for each experiment).

More than 400 vTRs have been identified in human viruses, predominantly in double-stranded DNA (dsDNA) viruses.^6^ Many vTRs are key mediators of viral pathogenesis, including well-characterized oncoproteins such as E1A from adenovirus, HBx from hepatitis B virus (HBV), and E6 and E7 from human papillomavirus (HPV).^14–16^ Other vTRs primarily function as immunosuppressive factors, such as vIRF1-3 and Rta from Kaposi’s Sarcoma-associated virus (KSHV).^17^ Notably, certain vTRs exhibit allele-specific binding, contributing to the pathogenesis of chronic or complex diseases. For example, the EBV EBNA2 is recruited to nearly half of the genetic risk loci associated with autoimmune disorders such as multiple sclerosis and systemic lupus erythematosus, where it shows evidence of allele-dependent.^18,19^

Despite their critical role in disease, mechanisms of host transcriptional modulation are poorly understood for most vTRs. For instance, genomic occupancy has been mapped for only 15 unique vTRs,^6,20,21^ and protein-protein interaction (PPI) data are available for just 74 of the ∼400 vTRs.^22^ Although recent transcriptional activity studies have been systematic, they have focused on 80 amino acid regions,^23^ potentially missing larger, synergistic, or context-dependent activation or repression domains. The lack of comprehensive studies comparing various molecular functions of vTRs across diverse viral families has significantly limited our understanding of their regulatory strategies and contributions to pathogenesis.

Here, we systematically characterize 95 vTRs from 31 human dsDNA viruses and retroviruses using a multimodal framework to define how vTRs reshape the human transcriptome and uncover the principles governing their regulatory functions. This vTR set spans different structural folds, timing of expression, reported functional roles, and likely mechanisms of action. We find that vTRs from phylogenetically distinct viruses often converge on core host pathways, including immunity, signal transduction, cell division, and apoptosis, revealing both shared and divergent regulatory strategies. This transcriptional rewiring is mediated by distinct mechanisms, including direct DNA binding (independently or cooperatively with hTFs), tethering to hTFs, and active disruption of hTF-DNA binding. We show that vTRs can function as transcriptional activators or repressors and engage in PPI networks that exhibit both shared and unique features compared to hTFs. This integrative analysis reveals previously unappreciated redundancies and division of labor among vTRs encoded by the same virus, as well as striking functional divergence among orthologs. Finally, we identify links between vTR-modulated genes and disease risk loci, suggesting that viral regulators may act as environmental sensitizers of genetic susceptibility. Collectively, our work highlights vTRs as a functionally diverse set of regulators and provides new insights into both viral pathogenesis and fundamental principles of transcriptional control.

## RESULTS

### vTR1.0: a resource to study the vTR functional landscape

Similar to human transcriptional regulators, vTRs function through direct or indirect recruitment to DNA regions and modulate gene expression through interactions with cofactors and hTFs. To determine the mechanisms by which different vTRs perturb gene regulatory networks, we used an extensive multimodal functional approach integrating transcriptomics, biophysical interactions, and measurements of transcriptional activity (**Figure 1B**). We selected a subset of 95 vTRs from the annotated compendium of 419 vTRs for gene synthesis (**Data S1**). These vTRs span all major dsDNA virus and retrovirus families (**Figure 1C**), and include several vTRs known to impact pathogenesis such as E6 and E7 from different HPV strains, EBNA2 from EBV, and E1A from adenovirus. Codon optimized vTR open reading frames were cloned into Gateway compatible donor vectors, resulting in the vTR1.0 entry clone collection, which are easily transferable to destination vectors for different functional assays (**Figures 1B-C**). To determine the effect of vTRs on the human transcriptome, we performed Bulk RNA Barcoding and sequencing (BRB-seq) ^24^ in HEK293T cells transiently expressing 84 different vTRs, of which 58 resulted in altered human gene expression using our stringent criteria (≥100 differentially expressed genes; |Log2FC| ≥ 0.5, p < 0.05). To evaluate the intrinsic ability of vTRs to bind to DNA regions (i.e., in the absence of other hTFs), we used enhanced yeast one-hybrid (eY1H) assays, a yeast-based reporter assay that detects protein-DNA interactions (PDIs) in a high-throughput manner. We tested the vTR1.0 collection against a set of 112 DNA-baits derived from the promoter elements (2 kb regions upstream of the transcription start site) of cytokine genes, which are heavily targeted by many viruses.^25,26^ To assess the possible co-binding or sequestration between hTFs and vTRs, we used paired yeast one-hybrid (pY1H) assays, a variation of eY1H that evaluates pairs of proteins for synergistic or antagonistic binding to DNA. In particular, we tested 113 vTR-hTF pairs against 39 immune gene promoters and 83 promoters of genes involved in cell signaling, tumor suppression, oncogenesis, differentiation, metabolism, replication, and apoptosis.

To determine the ability of vTRs to activate or repress transcription, we performed mammalian one-hybrid (M1H) assays in HEK293T cells, a method in which proteins are tethered to a firefly luciferase reporter construct to quantify transcriptional activation or repression. Finally, we used yeast two-hybrid (Y2H) assays to screen vTR1.0 against the entire human ORFeome v9.1, comprised of 17,472 protein-coding genes, and identified 132 PPIs between 33 vTRs and 67 human proteins. We maintained a similar viral family representation across assays relative to vTR1.0 (**Figure 1C**) and identified differentially expressed genes, intermolecular interactions, or significant transcriptional activity for 91% of vTRs tested (**Figure 1D** and **S1**). This illustrates the breadth of functional profiling data generated for our vTR compendium, which is necessary for deriving an integrated picture of vTR function and enabling systematic discovery and comparison of the molecular strategies viruses use to hijack human gene regulation.

### vTRs modulate multiple host pathways

To determine the pathways rewired by different vTRs, we used HEK293T cells as a standardized human background to allow systematic comparison across the vTRs profiled. This cell line provides high transfection efficiency and stable, well-characterized transcriptional programs, enabling robust detection of vTR-driven perturbations. Moreover, HEK293T cells express a broad set of transcriptional cofactors and signaling components, well-suited for other assays within our multimodal profiling. Importantly, their low basal antiviral response minimizes confounding immune activation, allowing us to quantify vTR-specific effects under basal and stimulated conditions. We expressed individual vTRs in HEK293T cells for 72 hs, determined the transcriptomic profile using BRB-seq, and identified the set of differentially expressed genes relative to empty vector. Gene set enrichment analysis of differentially expressed genes shows that vTRs generally dysregulate pathways associated with immune response, apoptosis, cell cycle, DNA damage repair, and metabolism (**Figure 2A**). These pathways have been broadly associated with viral replication and persistence, and some are known to be affected by vTRs; for example KSHV vIRFs are known to inhibit p53 activity.^27,28^ Consistent with this, vIRF1/3/4 downregulated several pathways associated with p53 regulation, as well as downstream pathways including those regulating cell cycle checkpoints and DNA double-stranded break repair (**Figure 2A**). Similarly, HPV E6 and E7, commonly associated with suppression of apoptosis and innate immunity and the upregulation of cell cycle genes,^29,30^ substantially modulated these pathways.

**Figure 2:**
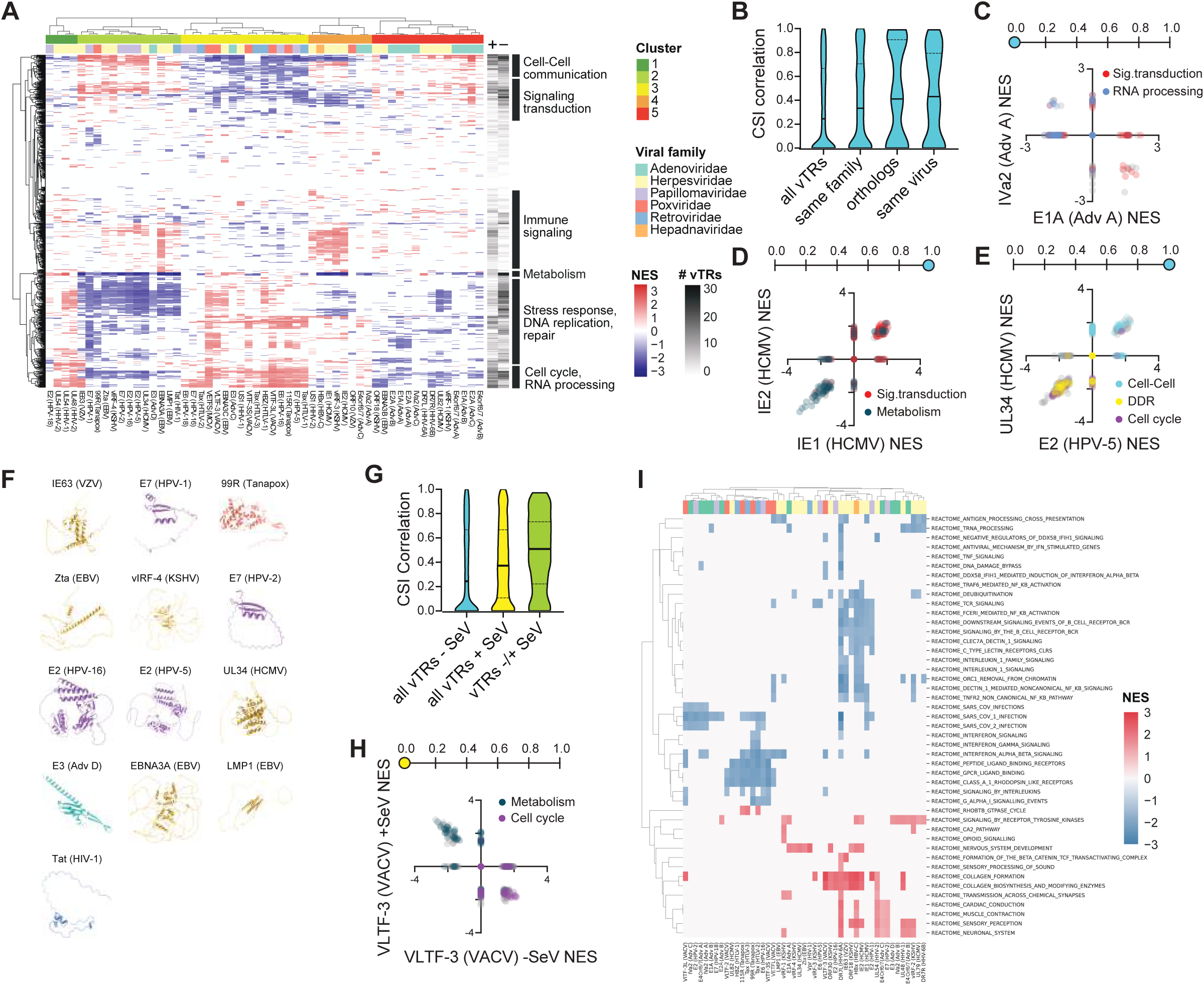
Perturbation of human gene expression by vTRs. (A) Aggregated results of gene set enrichment analysis from cells expressing different vTRs. vTR are clustered based on CSI-corrected Pearson correlation coefficient. Rows correspond to gene sets clustered based on gene similarity. Normalized enrichment scores (NES) are represented for each vTR as a color gradient. The number of vTRs with a positive or negative NES per pathway is indicated in grey. (B) Distribution of connection specificity index (CSI) for all pairs of vTRs, vTRs from the same family, vTR orthologs, and vTRs from the same virus. Solid lines denote the median and dashed lines denote quartiles. (C-E) Scatter plots comparing the NES of pathways dysregulated by pairs of vTRs. Number lines above indicate the CSI correlation for the given pair of vTRs. Selected pathways are colored. (C) Adenovirus A E1A versus IVa2. (D) HCMV IE1 versus IE2. (E) HPV-5 E2 versus HCMV UL34. (F) Alphafold2 models for vTRs from Cluster 2 in Figure 2A. Color indicates viral family. (G) Distribution of CSI correlation scores for all vTRs in resting cells, all vTRs tested in Sendai Virus-infected cells, and vTRs compared between untreated and Sendai Virus-infected cells. SeV - Sendai Virus. (H) Scatter plot comparing the NES of pathways dysregulated by vaccinia virus (VACV) VLTF-3 in untreated and Sendai Virus-infected cells. (I) Heatmap of NES from GSEA comparing SeV infection versus no infection in vector and vTR conditions. Pathways shown are significantly altered by SeV infection and display opposite responses between vector and vTR.

We next evaluated whether vTRs from the same viral family, vTR homologs from related viruses, and vTRs from the same virus broadly affect similar or different pathways. To accomplish this, we calculated the Pearson correlation coefficient of pathway normalized enrichment scores for each pair of vTRs, which were rescaled using the connection specificity index (CSI) that ranges from 0 (anti or not correlated) to 1 (highly correlated).^31,32^ We observed that vTR pairs have an overall low correlation between affected pathways, consistent with the diverse functions and pathways affected by different vTRs, whereas homologous vTRs and vTRs from the same virus have more similar transcriptional effects (**Figure 2B**). The broad distribution of correlation values between vTRs from the same virus, however, suggests that division of labor, redundancy, cooperation, and opposing effects between vTRs may all be employed by viruses to robustly rewire host cellular pathways. For instance, E1A and IVa2 from adenovirus A exhibit opposite regulation of signal transduction and RNA processing related pathways and mutually exclusive regulation of cell cycle (E1A) and metabolism (IVa2) (**Figure 2C** and **Data S2**). The contrasting transcriptional effects of these two vTRs combined with their distinct expression kinetics (E1A - immediate early, IVa2 - late) underscore the unique requirements for each progressive stage of adenovirus infection. Conversely, IE1 and IE2, both expressed from the human cytomegalovirus (HCMV) major immediate early promoter, have highly concordant effects on host pathways (**Figure 2D**), including upregulation of signal transduction and downregulation of metabolism pathways. Together, this illustrates how vTRs from the same virus may have similar or different transcriptional effects depending on their expression timing during infection.

Interestingly, we identified 70 vTR-pairs that produce highly similar transcriptional profiles (CSI > 0.9) from viruses belonging to different viral families. For example, HPV-5 E2 and HCMV UL34 (**Figure 2A**), upregulate pathways involved in cell-cell communication and downregulate those associated with cell cycle control and DNA damage repair (**Figure 2E**). This marked degree of pathway similarity between HPV-5 E2 and HCMV UL34, as well as other vTRs with similar transcriptional profiles, is not attributable to similarities in structure (**Figure 2F**). Together, these results illustrate how vTRs from distant, unrelated viruses can have similar transcriptional effects, suggesting that viruses have converged to affect overlapping host pathways while retaining specific effects on viral gene regulation.

More generally, we found 5 clusters of vTRs with overall similar transcriptional profiles (**Figure 2A**). As expected, homologous vTRs are generally clustered together, illustrating the quality of our clustering approach. Of the clusters identified, Clusters 2 and 3 are notable for exhibiting somewhat opposite effects. vTRs from Cluster 2 downregulate genes involved in cellular stress such as apoptosis, DNA damage response, hypoxia, and chemical stress, and those related to cell replication, and upregulate immune, cell-cell communication, and intracellular signaling pathways. Instead, vTRs in Cluster 3 include many oncoproteins (e.g, HPV-18 E6, HPV-16/HPV-5 E7, and HTLV-1 Tax^4,33^ and upregulate pathways related to cell cycle progression, DNA damage, and stress responses, while downregulating those involved in extracellular signaling and antigen presentation. Interestingly several viruses, such as HPV and EBV, encode vTRs present in both Clusters 2 and 3, suggesting potential balancing mechanisms controlling cell growth and stress at different stages in the viral replication cycle. In addition, each cluster is composed of vTRs from different viruses and families. Together, our results suggest that unrelated viruses have evolved convergent transcriptional strategies to manipulate common host processes, while retaining distinct programs tailored to their specific requirements for persistence and replication.

### vTR regulation of pathways in infected cells

Viral infection greatly affects cell states and signaling pathways, some of which are leveraged by the virus to promote replication or persistence, while others reflect host antiviral responses. To determine how vTRs function in infected cells, we performed BRB-seq in HEK293T cells infected with Sendai virus (SeV) following transient transfection of 49 vTRs. We found that vTRs often regulate similar pathways regardless of infection state (**Figure 2G**), suggesting that the function of most vTRs is robust to antiviral mechanisms. However, 12 vTRs affect different pathways in infected versus uninfected conditions, and in some cases have opposite effects. For instance, VLTF-3 (vaccinia virus) shifts from repressing to activating metabolic/biosynthetic pathways and from activating to repressing cell cycle pathways in SeV-infected cells (**Figure 2H**), which may enable the virus to adapt to different cellular states.

Several vTRs, including IE1 (HCMV) and KSHV vIRFs, are known to act as immunomodulatory proteins.^34–36^ To identify additional vTRs that may counteract the antiviral effects elicited by infection, we focused on 45 pathways that are altered under SeV infection relative to resting cells, including interferon response, interleukin signaling, NF-□B activation, and antigen presentation. We identified 47 vTRs that affect at least one of these pathways in SeV-infected cells (**Figure 2I**), of which 39 downregulate immune-associated pathways normally upregulated in the presence of SeV. While several of these associations are known, such as vIRF-1 affecting interferon pathways^36,37^ and IE1/2 downregulating the T-cell receptor and interleukins,^34,35^ our results also identify several novel immunoregulatory functions. For example, Tax proteins from HTLV-2 and HTLV-3, typically associated with NF-□B regulation,^38^ both downregulate type I interferon signaling, whereas DR7L (HHV-6A) and IE63 (Varicella-Zoster virus - VZV) downregulate IL-1-associated pathways (**Figure 2I**). In addition to immunomodulatory roles, 31 vTRs upregulate signaling pathways normally suppressed during active infection (**Figure 2I**), which may reflect indirect immune evasion strategies that attempt to reestablish cellular processes needed for replication. Together, these findings reveal that vTRs not only buffer against infection-induced stresses but also actively reshape antiviral signaling.

### vTRs are recruited to host DNA through multiple mechanisms

Transcriptional regulators can be recruited to DNA elements using different mechanisms, including direct binding to DNA (either independently or as part of a protein complex) and indirect recruitment mediated by DNA-binding factors. We previously curated 33 ChIP-seq data sets corresponding to 12 vTRs, which showed large variations in target preference and binding distance to transcription start sites.^6^ This study showed that ChIP-seq peaks of many vTRs are enriched for hTF motifs, suggesting indirect recruitment of vTRs to genomic sites. Other vTRs have been shown to bind DNA directly, including bZIP Zta (EBV) and E2A (adenovirus).^12,39^

To identify which vTRs are recruited to DNA directly and through which mechanisms, we first used PADIT-seq^40^ to assay the DNA binding activities of ten vTRs to all possible 10-bp DNA sequences. These vTRs were selected based on their likelihood of directly binding to DNA given domain presence (e.g, viral IRFs), Y1H data, or previous vTR-DNA interaction data.^26^ To facilitate protein expression, we expressed structured vTR regions and 10-15 flanking amino acids, as previously conducted for hTFs.^40^ However, none of these vTR constructs showed significant binding to 10-bp DNA sequences, while our hTF positive controls did. The observed lack of DNA binding activity may be due to misfolding of the vTRs in the *in vitro* transcription and translation system, the need for excluded vTR regions, or interference by the ALFA tag fusion required by the assay, that these vTRs can only bind sequences longer than the 10-mers tested in PADIT-seq, or that the vTRs associate indirectly, rather than directly, with their target genomic sites.

To address these possibilities, we performed eY1H assays which evaluate interactions between individual proteins and DNA regions of up to 2 kb in the absence of other human or viral proteins. We tested 95 vTRs against 112 immune gene promoters, given that most vTRs dysregulate at least one immune pathway. We also assessed cooperativity with hTFs using pY1H assays, a modified eY1H approach that evaluates DNA binding of protein pairs. We tested 113 vTR-hTF pairs selected based on reported PPIs, homologs of known vTR-hTF interactions, and PPIs identified in this study. Both individual vTRs/hTFs and their pairs were assayed against 39 cytokine promoters and 83 promoters of genes from the Cancer Gene Census, involved in cell proliferation, cell death, and cell signaling, all pathways dysregulated by vTRs (**Figure 2A**).

We identified 17 vTRs (18% of vTRs tested) that bind human promoters (**Figure 3A** and **Data S3** and **S4**). This is similar to the proportion of hTFs binding these regions (24%),^25^ suggesting that many vTRs might not rely on hTFs for DNA recruitment, in contrast to previous studies.^6^ Vaccinia VLTF-1 and hepatitis B X-protein (HBx) have the most DNA targets, with 39 and 14 PDIs respectively. In addition to the established HBx target *TNF*,^41^ HBx binds the *MYC* promoter, suggesting direct regulation, consistent with reports of HBx-associated increases in *MYC* expression and activity in hepatocellular carcinoma, though direct control had not been shown.^42,43^ We also found HBx binding to the promoter of *LEF1*, a key hTF controlling Wnt signaling, in line with our transcriptomics data showing HBx upregulates several Wnt signaling pathways and with previous studies that implicate HBx in regulating host Wnt signaling during the progression of hepatocellular carcinoma.^44,45^

**Figure 3:**
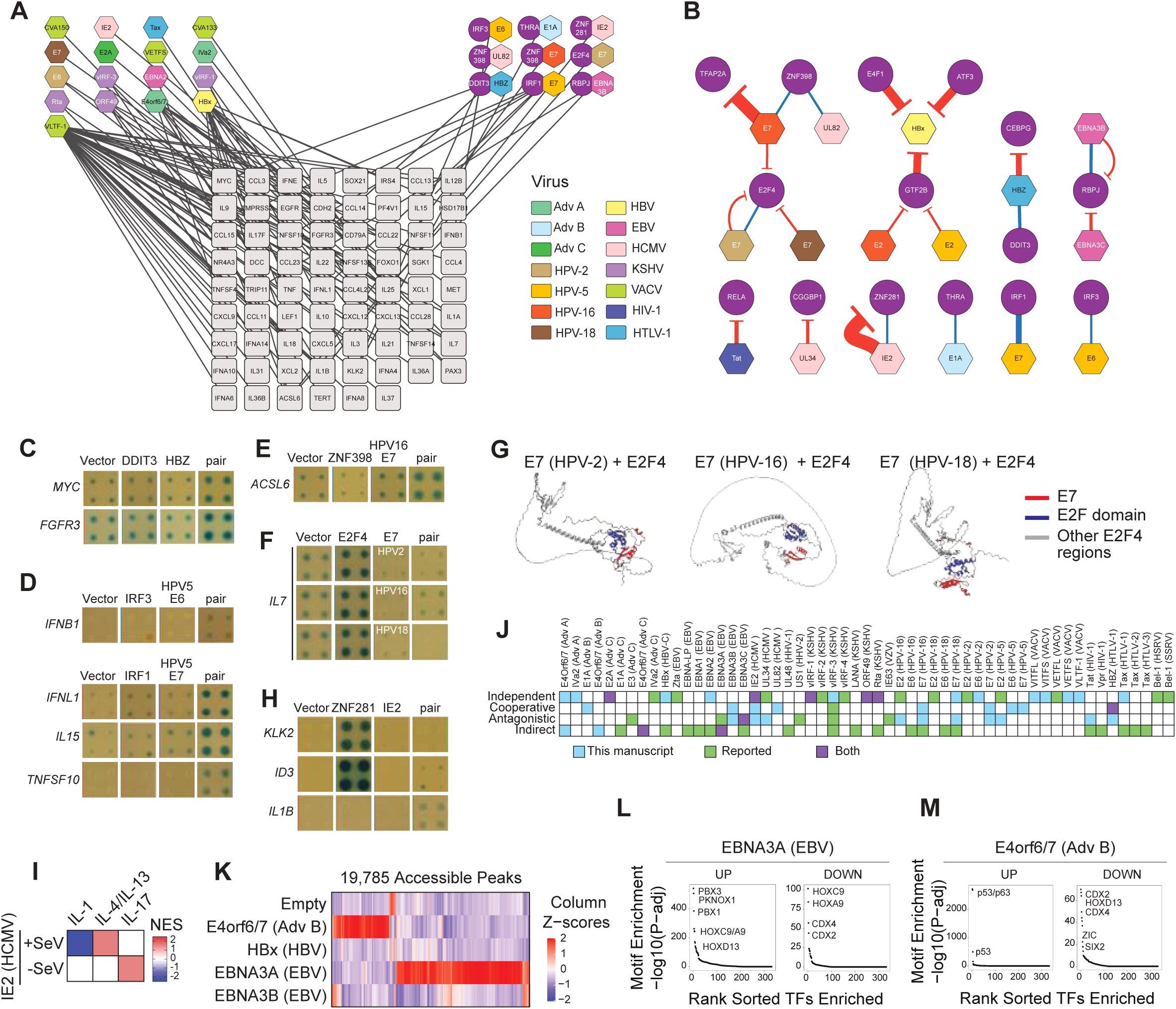
vTRs bind host regulatory elements and alter chromatin structure. (A) Protein-DNA interactions for vTRs (left), vTRs cooperating with hTFs (right), and human promoter sequences. Hexagons represent vTRs, and are colored by virus. Purple circles represent hTFs. Squares represent human promoters. (B) Network of relationships between hTFs and vTRs. Blue edges represent cooperativity, red edges represent antagonism. Edge thickness indicates the number of promoters at which a specific interaction occurs. (C–F and H) Representative pY1H interactions for hTF-vTR pairs. Italicized gene names (left) indicate the tested promoter bait. Yeast strains shown: empty vector control, hTF only, vTR only, hTF-vTR pair. (C) Cooperativity between DDIT3 and HBZ (HTLV-1) at MYC and FGFR3 promoters. (D) Cooperativity between IRF3 and E6 (HPV-5) at IFNB1 promoter. Cooperativity between IRF1 and E7 (HPV-5) at IFNL1, IL15, and TNFSF10 promoters. (E) Cooperativity between ZNF398 and E7 (HPV-16) at ACSL6 promoter. (F) Antagonism of E2F4 by multiple E7 homologs at the IL7 promoter. (H) Antagonism of ZNF281 by IE2 (HCMV) at KLK2 and ID3 promoters; cooperativity between ZNF281 and IE2 at IL1B promoter. (G) Alphafold3 models of interactions between E2F4 and E7 from different HPV strains. (I) Heatmap of NES scores for Interleukin pathways derived from BRB-seq data for IE2 (HCMV) in SeV-infected and uninfected cells. (J) vTR DNA binding modalities described in this manuscript (light blue), reported in the literature (green), or both (purple). (K) Heatmap of z-score normalized chromatin accessibility at 19,785 peaks that are differentially accessible (FDR ≤ 0.1 and log2FC ≥ 1) in at least one vTR surveyed via PROD-ATAC comparing E4ORF6/7 (Adv B), HBx (HBV), EBNA3A (EBV), and EBNA3B (EBV). Columns of peaks are hierarchically clustered. (L-M) Representative DNA motif enrichment within increased/decreased differentially accessible peaks for EBV EBNA3A (L) and Adenovirus B E4ORF6/7 (M). Displayed values were calculated using hypergeometric tests and plotted relative to the ranked order of all queried motifs.

We also identified 9 vTRs that bind DNA cooperatively with hTFs (**Figure 3A-B**). For example, we observed co-binding between bZIP HBZ (HTLV-1) and the human bZIP DDIT3 at the *MYC* and *FGFR3* promoters (**Figures 3C**), two genes heavily involved in oncogenic pathways. This is consistent with studies showing that HTLV-1 activates *MYC* expression in adult T-cell leukemias.^46^ We also found cooperative interactions between HPV-5 E6 and IRF3 at the *IFNB1* promoter, and between HPV-5 E7 and IRF1 at the *IFNL1*, *IL15*, and *TNFSF10* promoters (**Figure 3D**), in line with known immunomodulatory roles of E6 and E7.^47–49^ Considering that E6 and E7 are known to inhibit transcriptional activation by IRF3 and IRF1, respectively,^49,50^ our results suggest that E6 and E7 may outcompete IRF dimers to prevent activation of immune response genes. Interestingly, HPV-16 E7 cooperatively binds to DNA with ZNF398 (**Figure 3E**), an oncogenic hTF that promotes tumorigenesis through p53 degradation, enhanced cellular metabolism, disruption of ferroptosis, and maintenance of cancer stem cell niches.^51–55^ Our results suggest that E7 may leverage these ZNF398 functions to promote cell proliferation and prevent cell death.

In addition to the detected vTR binding events, we observed 12 vTR-hTF pairs in which a vTR antagonized the binding of a hTF at specific promoters (**Figure 3B**). This likely occurs by sequestration, as Alphafold3 modeling predicts binding of the vTRs to the DNA binding domain of the interacting hTFs (**Figure S2**). For example, E2F4 is antagonized by several HPV E7 orthologs, which are predicted to interact with the DNA-contacting interfase of the E2F domain (**Figure 3B, F, G**). Ingenuity pathway analysis of our BRB-seq data shows that expression of HPV-16 E7 leads to downregulation of E2F targets (z-score = −2.164, p = 0.0022). HPV E7 is known to bind the tumor suppressor retinoblastoma gene (RB1) and prevent its association with E2F proteins.^56,57^ Together with our results, this suggests that E7 may disrupt E2F function through two separate mechanisms: inhibition of RB1-E2F association and sequestration of E2F, preventing its association with DNA. We observe a similar pattern for TFAP2A, which was antagonized by HPV-16 E7 at 15 promoters (**Figure 3B**) and was previously shown to be downregulated in cervical cancer cell lines expressing E7.^58^

We also observed 36 instances in which HCMV IE2 antagonized the DNA binding of ZNF281 likely by forming an interlocked complex between both proteins that occludes the cys2His2 zinc fingers of the DNA binding domain (**Figures 3B, 3H, S2,** and **S3**). ZNF281 is an oncogenic hTF associated with several types of cancer and plays a role in inflammation.^59–62^ By preventing ZNF281 binding to its normal targets, IE2 may redirect ZNF281 either to the HCMV genome or to other host gene promoters. Interestingly, we detected a cooperative interaction between IE2 and ZNF281 at the *IL1B* promoter (**Figures 3B** and **3G**). Although IE2 is primarily an activator, we propose that recruitment of ZNF281 may result in a repressive complex. Indeed, we observed that IE2 repressed IL1-related pathways in our transcriptomics data from SeV-infected cells (**Figure 3I**). Our results suggest a novel regulatory mechanism by which IE2 co-opts ZNF281 to suppress interleukin response during active infection states. This mechanism is distinct and complementary to reported immune suppressor activities of IE2 mediating STING and CD83 degradation, activation of the IL10/STAT3 signaling pathway, and inhibition of antigen presentation.^63–66^

Together, our results extend previous reports on vTR recruitment to host DNA and show that in addition to indirect and independent DNA binding, vTRs may reprogram host gene expression through cooperative or antagonistic interactions with hTFs (**Figure 3J**). Interestingly, we show that 18 vTRs may modulate expression through multiple mechanisms, potentially expanding their regulatory targets (**Figure 3J**). This is likely an underestimate given the limited number of DNA sequences tested by Y1H assays and the sparsity in currently available vTR ChIP-seq datasets. Altogether, our data uncover an unexpectedly broad repertoire of DNA-engagement mechanisms, showing that vTRs can operate as independent DNA binders, partners that co-opt hTFs, or antagonists that block hTF interactions, often combining multiple modes to rewire host gene expression.

### vTRs alter chromatin accessibility

In addition to binding to open regions in the host genome, vTRs can drive changes in chromatin architecture. While these effects have been demonstrated for several vTRs including Adenovirus E1A, EBV BZLF1, and HIV Tat,^67–69^ the effects of many vTRs on chromatin accessibility remain largely uncharacterized. We performed PROD-ATAC, a single-cell ATAC-seq approach that evaluates the effect of protein overexpression on chromatin accessibility (39048711), to determine the accessibility landscapes generated by EBV EBNA3A and EBNA3B, Adenovirus B E4ORF6/7, and HBV X-protein expression in HEK293T cells (**Figure S4A**). Of these vTRs, EBNA3A produced the largest number of differentially accessible peaks (**Figure 3K**), with newly opened regions enriched for PBX, CDX, and HOX motifs (**Figure 3L**). This is consistent with a recent study showing that EBNA3A ChIP-seq peaks and newly opened regions (ATAC-seq) are enriched for PBX motifs.^20^ We also found that regions closed in EBNA3A expressing cells are enriched for HOX and CDX motifs, though these regions are different from those with HOX/CDX motifs opened in EBNA3A-expressing cells, suggesting that EBNA3A induces the remodeling of chromatin occupied by different homeodomain proteins.

We also detected marked changes in accessibility by Adenovirus B E4ORF6/7, which exhibit minimal overlap with those driven by EBNA3A (**Figure 3K**). Newly opened chromatin induced by E4ORF6/7 was enriched for p53/p63 motifs (**Figure 3M**), whereas closed regions were dominated by HOX/CDX motifs. E4ORF6/7 has been reported to interact with p53; however, unlike its larger splice variant E4ORF6, it does not degrade p53.^70,71^ Together with our results, this supports a model in which E4ORF6/7 promotes occupancy of p53 binding sites, potentially modulating p53 targets. Indeed, ingenuity pathway analysis shows that Adenovirus B E4ORF6/7 expression leads to the upregulation of p53-responsive genes (z-score = 2.092, p = 4.2×10-5). Interestingly, both E4 splice variants are co-expressed early during infection, although E4ORF6/7 is less stable,^72^ suggesting that the adenoviral lifecycle may require distinct regulation of p53 (e.g. degradation/transcriptional regulation) at specific time points during infection.

### vTRs have intrinsic transcriptional effector activity

Like hTFs, many vTRs contain effector domains that recruit transcriptional cofactors to activate or repress target gene expression.^73^ A previous study using HT-recruit, a high-throughput tiling peptide approach to screen for transcriptional activity, identified hundreds of viral effector domains.^23^ Because effector domains may function differently in the context of a full-length protein structure, and because some effector domains can be larger than those examined using HT-recruit (80 amino acids),^74,75^ we interrogated the transcriptional activity of 95 full-length vTRs using M1H assays. M1H uses a luciferase reporter downstream of an enhancer element that contains 4 copies of the Gal4 upstream activating sequence (UAS). A protein (e.g., a full-length vTR) fused to the Gal4 DNA binding domain is then recruited to the reporter construct and may activate or repress luciferase expression. We used two versions of the M1H assay, using either a minimal or a moderate promoter between the UAS and luciferase, to better capture both activation and repression. Using our minimal promoter system, we observed that 25.2% of vTRs tested are strong transcriptional activators (Log2(fold change) ≥ 5), compared to only 6.75% for a set 224 hTFs with broad representation across families and that was evaluated using the same M1H system (**Figure 4A**).^76^ This suggests that vTRs may have evolved potent effector functions, enabling viruses to more effectively reprogram viral and host transcriptional networks.

**Figure 4:**
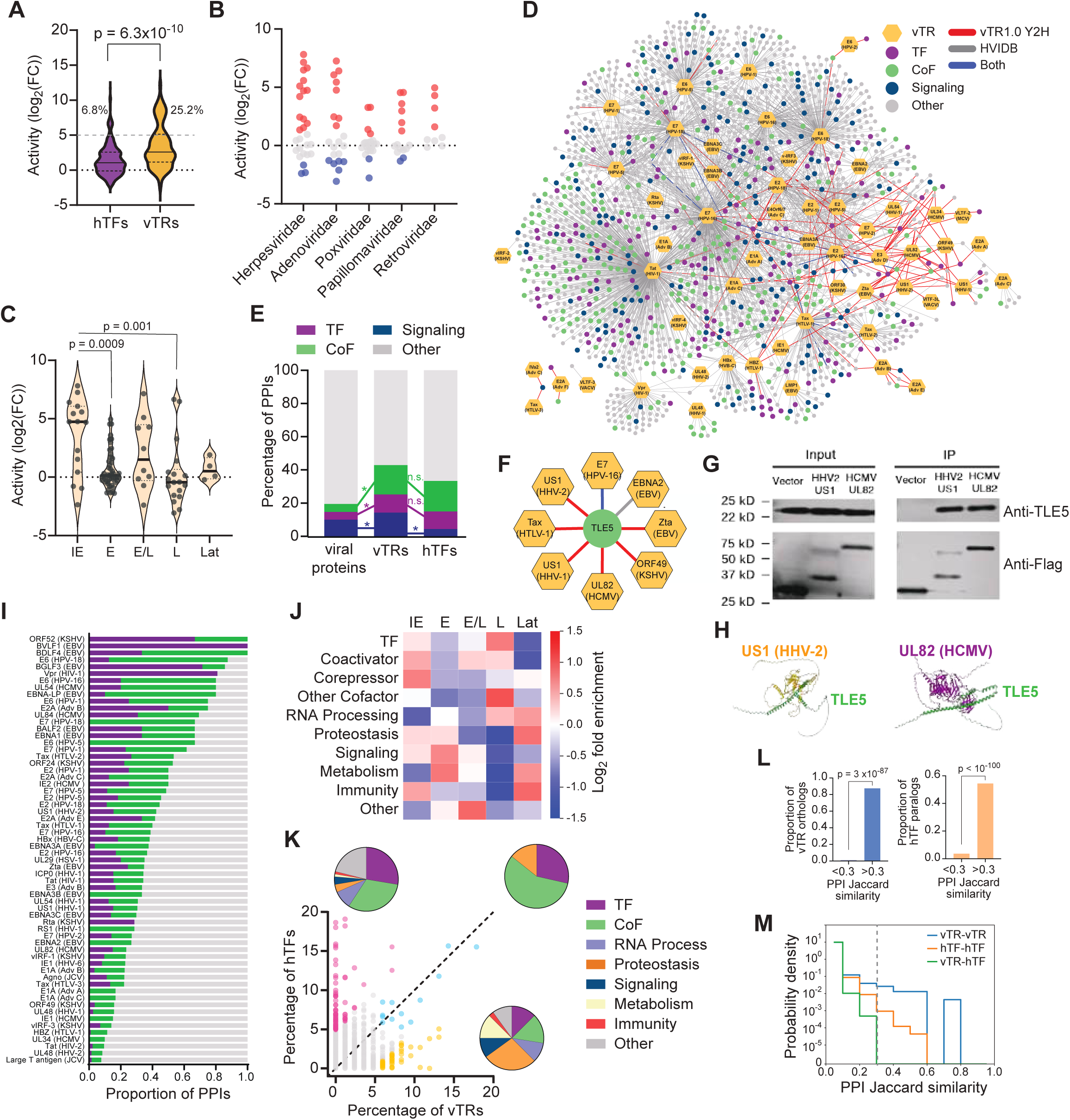
vTRs display intrinsic transcription activity and interact with host regulators. (A) Distribution of transcriptional activities elicited by hTFs and vTRs by M1H assays using a minimal promoter. Significance determined by two-tailed Mann-Whitney test. (B) Transcriptional activities of vTRs across viral families by M1H assays with a moderate promoter. Red and blue points indicate activating (L2FC ≥ 1) and repressing (L2FC ≤ −1) vTRs, respectively. (C) Distribution of transcriptional activities for vTRs expressed at different timepoints of infection. IE, immediate early; E, early; E/L, early/late; L, late; Lat, latent. Distributions were compared using One-Way ANOVA. Asterisks correspond to multiple comparisons adjusted p-values (*** P < 0.001). (D) Protein-protein interaction (PPI) network curated from HVIDB (http://zzdlab.com/hvidb) and the Y2H screen. Hexagons indicate vTRs and circles indicate human proteins. Edges indicate the source of the PPI. hTF, human transcription factor; CoF, cofactor. (E) Proportion of PPIs between viral proteins, vTRs, or hTFs and hTFs, cofactors (CoF), signaling proteins, or other proteins. *p<0.01 by two-tailed proportion comparison test. (F) PPI Sub-network between 8 vTRs and TLE5. (G) Western blot showing co-immunoprecipitation of TLE5 with FLAG-tagged HHV2 US1 and HCMV UL82. (H) Alphafold3 model of interactions between TLE5 and HHV2 US1 and HCMV UL82. (I) Proportion of PPIs between vTRs and hTFs, CoFs or other proteins. Only vTRs with at least 3 human PPIs are shown. (J) PPI log2fold enrichment between vTRs expressed at different timepoints of infection for ten categories of human proteins. (K) Percentage of vTRs or hTFs with PPIs with individual human proteins. Interactions shown are the union of vTR1.0 Y2H, HuRI (http://www.interactome-atlas.org), HVIDB, and BioGRID. Magenta dots indicate human proteins preferentially interacting with hTFs, gold dots indicate proteins preferentially interacting with vTRs, and cyan dots indicate proteins that frequently interact with both vTRs and hTFs. Pie charts represent the fraction of different protein classes within each previously described subset of proteins. (L) Proportion of vTR homologs (left) or hTF paralogs (right) represented in comparisons with high jaccard similarity (>0.3) or low jaccard similarity (<0.3). (M) Probability density distributions of Jaccard PPI profile similarity scores within vTRs (blue), within hTFs (orange), or between vTRs and hTFs (green).

Using our moderate promoter M1H system, we identified 36 activator and 13 repressor vTRs (**Figure 4B** and **Data S5**). We observed different distributions in vTR activity across viral families. Many highly active vTRs belong to herpesviruses or adenoviruses, the latter of which also account for half of repressive vTRs. Comparatively, poxvirus, papillomavirus, and retrovirus vTRs show milder transcriptional effects, which may reflect different roles in direct regulation of transcription.

We next compared the transcriptional activity of vTRs expressed at different stages of the viral lifecycle, and observed striking differences in activity between these groups (**Figure 4C**). Immediate-early vTRs, which play integral roles initiating viral expression cascades, have the highest activity among all groups. We also observed that many vTRs expressed late in the lytic cycle act as repressors rather than activators. Notable exceptions are highly active tegument proteins UL48 (HHV-1/2) which, although lately expressed, initiate transcription of immediate-early genes upon infection or reactivation (**Figure 4C**).^77^ Finally, vTRs expressed during latency exhibit low levels of transcriptional activity, suggesting that these vTRs may elicit more subtle or context-specific effects on viral and host expression to favor persistence.

### vTRs interact with overlapping but distinct sets of proteins compared to hTFs

hTFs regulate gene expression through interactions with cofactors, chromatin modifying enzymes, other hTFs, and members of the transcription preinitiation complex.^78–82^ To identify potential mechanistic similarities and differences between vTRs and hTFs, we compared their PPI profiles. Many PPIs between viral and host proteins were previously determined,^22,83,84^ mostly using affinity-based methods such as IP-MS; however, these methods are biased towards abundant proteins and often identify indirect interactors. Because we expected vTRs to interact with hTFs, which are generally lowly abundant, we performed Y2H assays, which do not depend on *in vivo* protein abundance as proteins are expressed using a defined promoter. We performed Y2H assays between each vTR in the vTR1.0 collection and the complete human ORFeome (∼17,500 proteins). Positive hits were retested pairwise in a matrix format to enable PPI comparisons between vTRs. In total, we identified 132 PPIs between 33 vTRs and 67 human proteins, 13 of which had been identified in earlier studies (**Figure 1D** and **Data S6**).

After integrating our Y2H with available PPI data, we observed that 31.2% of vTR interactions involve hTFs or cofactors, significantly more than for other viral proteins and similar to hTFs (**Figure 4D-E**). Strikingly, we detected previously unreported interactions between six vTRs and the cofactor TLE5 via Y2H assays (**Figure 4F**). TLE5 belongs to the TLE family of transcriptional co-repressors, which regulate proliferation, survival, cell fate, and immunity.^85,86^ Unlike other TLE proteins, TLE5 lacks repression domains and instead acts as a dominant-negative regulator interfering with repressive complex formation.^85–87^ We found that US1 (HHV-1), UL83 (HCMV), E7 (HPV-16), and Tax (HTLV-1), with available BRB-seq data, altered pathways typically regulated by TLE proteins, including Notch and Wnt signaling, and RUNX target genes (**Data S2**). We confirmed TLE5 interactions with US1 (HSV-2) and UL82 (HCMV) by co-immunoprecipitation and western blot (**Figure 4G**). Structural modeling of these interactions with Alphafold3 showed that both US1 and UL82 associate with the Q-domain of TLE5 (**Figure 4H**), which is required for interaction with other TLE proteins.^86,87^ This suggests that many vTRs from different viral families sequester TLE5 away from other TLE family members, thereby disrupting host transcriptional repression. Beyond TLE5, many other vTRs also modulate Notch and Wnt signaling pathways likely through alternative mechanisms, implicating these pathways as critical for viral replication or immune evasion (**Figure 2A** and **Data S2**).

Strikingly, over 60% of vTRs with at least 3 PPIs predominantly interact with non-TF/cofactor proteins (**Figure 4I**). This suggests a spectrum of vTR functionality beyond direct transcriptional regulation, consistent with the known multifunctionality of viral proteins including vTRs.^6^ For instance, early vTRs interact with proteins involved in proteostasis, signaling, and metabolism, aligned with their role in manipulating the host cell environment for viral replication (**Figure 4J**). Latent vTRs interact with immune, RNA processing, metabolic, and proteostatic proteins, potentially contributing to regulate cellular states, evade antiviral responses, and preserve latency.

Next, we evaluated which individual human proteins preferentially interact with vTRs or hTFs. As expected, many hTFs, cofactors, and RNA processing proteins preferentially interact with hTFs (**Figure 4K**). Similarly, a set of 14 proteins, consisting primarily of broadly expressed cofactors (e.g., EP300, CREBBP, and KAT2B) and hTFs (TP53, SP1, and CEBPB), was found to frequently interact with both hTFs and vTRs. We also found 40 proteins biased towards interactions with vTRs. This includes basal hTFs such as TBP and GTF2B, MEOX1 known to influence development, and NR4A1, RB1 and CCNA2, involved cell cycle control and cell death.^88–94^ We also found that proteins involved in proteostasis, cell signaling, metabolism, and immunity preferentially interact with vTRs rather than hTFs (**Figure 4K**), consistent with the multifunctionality of vTRs.^6^

Proteins with similar PPIs often share molecular functions. We therefore examined whether any vTR would engage in similar PPIs to those of a hTF, which could indicate convergent regulatory strategies. As expected, we observed enrichment of vTR homologs and hTF paralogs among vTR-vTR and hTF-hTF pairs with similar PPIs (**Figure 4L**); however, hTFs and vTRs engage broadly distinct sets of host proteins and we detected no hTF-vTR pair with similar PPIs (Jaccard>0.3, **Figure 4M**). This underscores that vTRs have evolved unique interaction networks rather than mirroring those of hTFs.

### Redundancy and division of labor between vTRs from the same virus

Large dsDNA viruses such as herpesviruses encode multiple vTRs that regulate diverse viral and host processes.^6^ To better understand the division of labor between vTRs from the same virus, we compared the overall host transcriptional effects of vTRs from viruses with two or more vTRs in our BRB-seq dataset. vTRs from the same virus range from affecting highly similar to highly dissimilar pathways under both resting and SeV-infected conditions (**Figures 5A-B**). Similarity in pathways affected is not necessarily related to timing of vTR expression (**Figure 5C**). For example, LMP1 and EBNA3A from EBV, both expressed primarily during latency, affect similar pathways to the lytic gene BZLF1, mostly driven by cell cycle, DNA damage, and immune/stress pathways (**Figure 5D-E**). By contrast, EBNA3A, EBNA3B, and EBNA3C, which are generated by alternative splicing from a single latently expressed transcript,^95^ exert distinct effects on host gene expression (**Figures 5D-E**). In contrast to EBV, HCMV vTRs expressed at the same time during infection exhibit the most correlated transcriptional effects (**Figure 5F**). For instance, immediate early IE1 and IE2 affect similar cell cycle, DNA damage, metabolism, and immune/stress pathways, whereas early UL34 and UL82 regulate cell cycle and immune/stress genes (**Figures 2D** and **5G**). Together, this suggests that vTRs from the same virus often co-regulate pathways, which is not necessarily related to the timing of vTR expression. These observations are consistent with a recent study showing that latent EBNA2 and lytic switch Rta from EBV share many gene targets during distinct stages of infection.^20^

**Figure 5:**
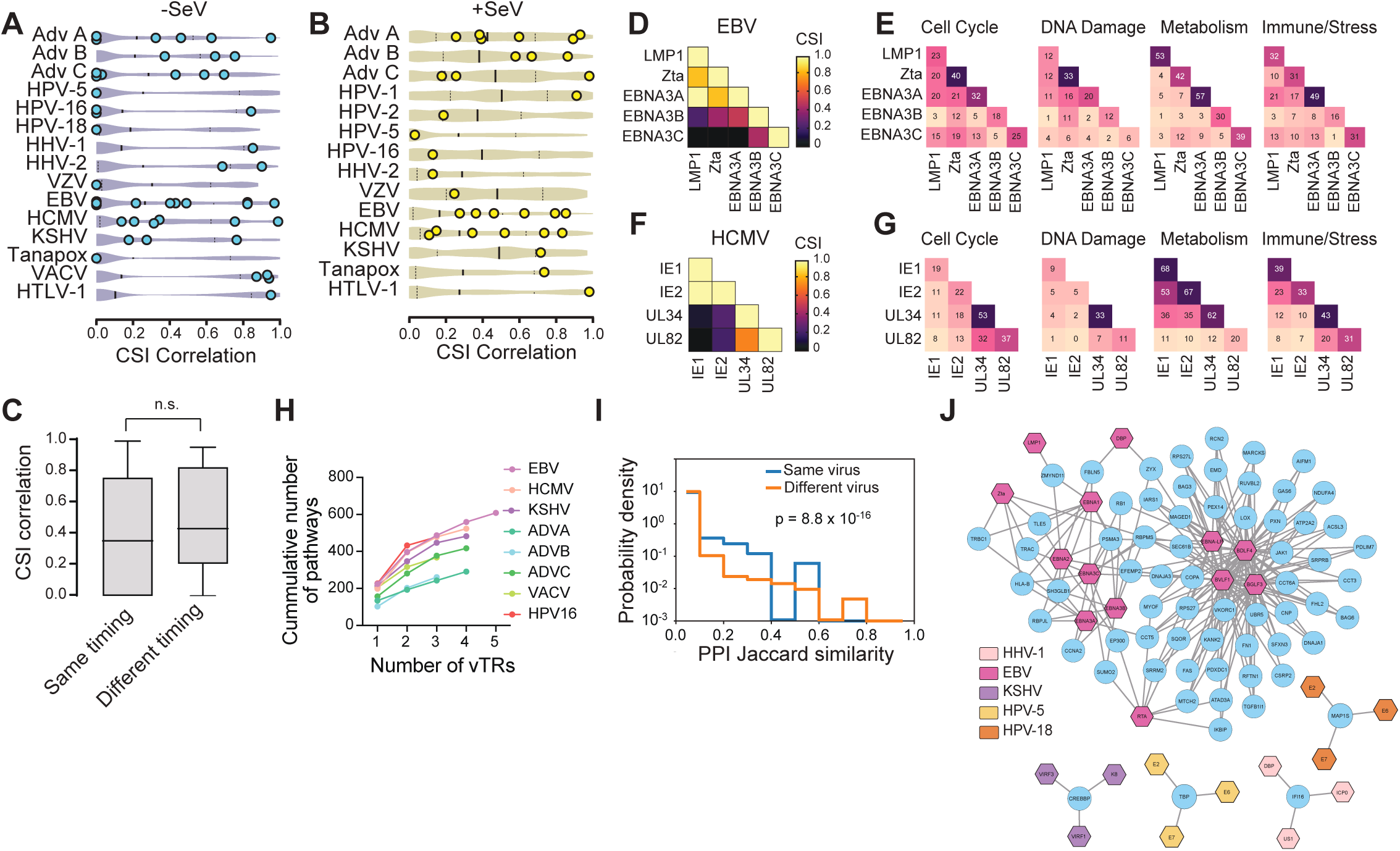
Functional diversity and redundancy between vTRs from the same virus. (A–B) CSI correlation scores for pairs of vTRs encoded by the same virus in resting (A) and SeV-infected (B) cells. For each virus, points represent within-virus vTR comparisons, shown relative to the distribution of all other possible pairwise comparisons involving these vTRs. (C) Distributions of CSI correlation scores for vTR pairs from the same virus with either the same or different expression timing. (D, F) CSI correlation score comparisons for five EBV (D) and four HCMV (F) vTRs in resting cells. (E, G) Heatmaps showing the number of overlapping pathways regulated by EBV vTRs (E) or HCMV vTRs (G) across four functional categories. Diagonal cells indicate the total pathways regulated by each vTR, and off-diagonal cells indicate the number of pathways shared between vTR pairs. (H) Cumulative number of pathways affected by the vTR repertoire of different viruses. Only viruses with at least 3 vTRs in our dataset are shown. (I) Probability density distributions of jaccard PPI profile similarity scores for vTRs from the same virus (blue) or different viruses (orange). Significance determined by two-sided Mann-Whitney’s U-test. (J) PPI networks for each virus showing human proteins that interact with at least three vTRs. Hexagons indicate vTRs and circles indicate human proteins.

vTRs from the same virus can also engage in division of labor, with individual vTRs preferentially influencing specific host pathways or exerting distinct effects across cellular conditions. For instance, while LMP1, Zta, and EBNA3A affect similar cell cycle, DNA damage, and immune/stress pathways, they differ in the particular metabolic pathways affected (**Figure 5E**). Other vTRs, instead, modulate shared or distinct pathways depending on particular cellular conditions. For example, UL48 and UL54 (HHV-2), which are highly concordant in non-infected cells (CSI = 0.9) diverge under SeV infection (CSI = 0.13), whereas 99R and VITF-3 (Tanapox) affect different pathways in non-infected conditions, but similar pathways under SeV infection (CSI = 0 uninfected cells; CSI = 0.73 infected cells) (**Figure 5B**). Together, this illustrates both partial redundancy and specialization between vTRs from the same virus in rewiring host gene regulation networks, which overall results in an expanded set of pathways affected (**Figure 5H**).

### vTRs from the same virus interact with similar host proteins

We next evaluated whether the overall shared pathways between vTRs from the same virus (**Figure 2B**) can be attributed to similar interactions with host proteins. We observed that vTRs from the same virus tend to have higher interaction profile similarity than vTRs from different viruses, suggesting partial overlap in their regulatory targets or reliance on similar host pathways during infection (**Figure 5I**). To explore how multiple vTRs from a single virus may act in concert, we examined host proteins interacting with at least three vTRs from the same virus in our dataset (**Figure 5J**). In line with the overall enrichment of transcriptional cofactors among vTR targets, KSHV and EBV vTRs interact with CREBBP and EP300, whereas HPV-5 vTRs bind to TBP. Beyond transcriptional regulators, we identified shared interactions involving diverse cellular processes: HHV-1 vTRs associate with the immune sensor IFI16, HPV-18 vTRs engage the cytoskeletal and autophagy-related protein MAP1S, and EBV vTRs collectively interact with host factors linked to signaling, apoptosis, immune responses, and metabolism (**Figure 5J**). These observations indicate that vTRs from the same virus can converge on common host interaction hubs, suggesting coordinated modulation of cellular pathways during infection.

### vTR homologs display varying levels of functional divergence

Homologous viral genes range from functionally conserved to highly divergent, reflecting selective pressures for either conservation or adaptation to specific tropisms and hosts. We compared 19 sets of vTR homologs across our experimental modalities (**Figure 6A**). Despite an overall high correlation in transcriptional effects between homologous vTRs relative to random vTR-pairs in our BRB-seq dataset (**Figure 2B**), homologs display varying levels of similarity (**Figure 6B**). For instance, HTLV Tax proteins are highly correlated with each other, sharing many regulated pathways in both uninfected and infected conditions. By contrast ICP22 homologs from alpha-herpesviruses (HHV-1 US1, HHV-2 US1, and VZV IE63) affect different pathways, with VZV IE63 being the most different, consistent with the evolutionary distance between VZV and HHV-1/2. In particular, HHV-1 and HHV-2 US1 both downregulate similar metabolic pathways, while these are upregulated by VZV IE63. All three ICP22 homologs, however, have very different effects on cell-cell signaling pathways (**Figure 6C**). While HHV-1 US1 mostly downregulates pathways associated with extracellular signaling, HHV-2 US1 and VZV IE63 frequently upregulate them; however, the identity of these upregulated pathways generally differs (**Figure 6C**).^96^ The pathways exclusive to HHV-2 US1 are largely involved in cell growth and metabolite trafficking, whereas VZV IE63 upregulates pathways associated with extracellular matrix organization and interaction between cells. Unlike HHV-1/2, where transmission is mediated by virion release and entry, VZV is mostly transmitted through cell-to-cell contact.^97,98^ Thus, targeted regulation of host pathways facilitating cell-cell contact by VZV IE63 and not other alpha-herpesvirus ICP22 homologs underscores a major difference in pathology and transmission of these three viral species.

**Figure 6:**
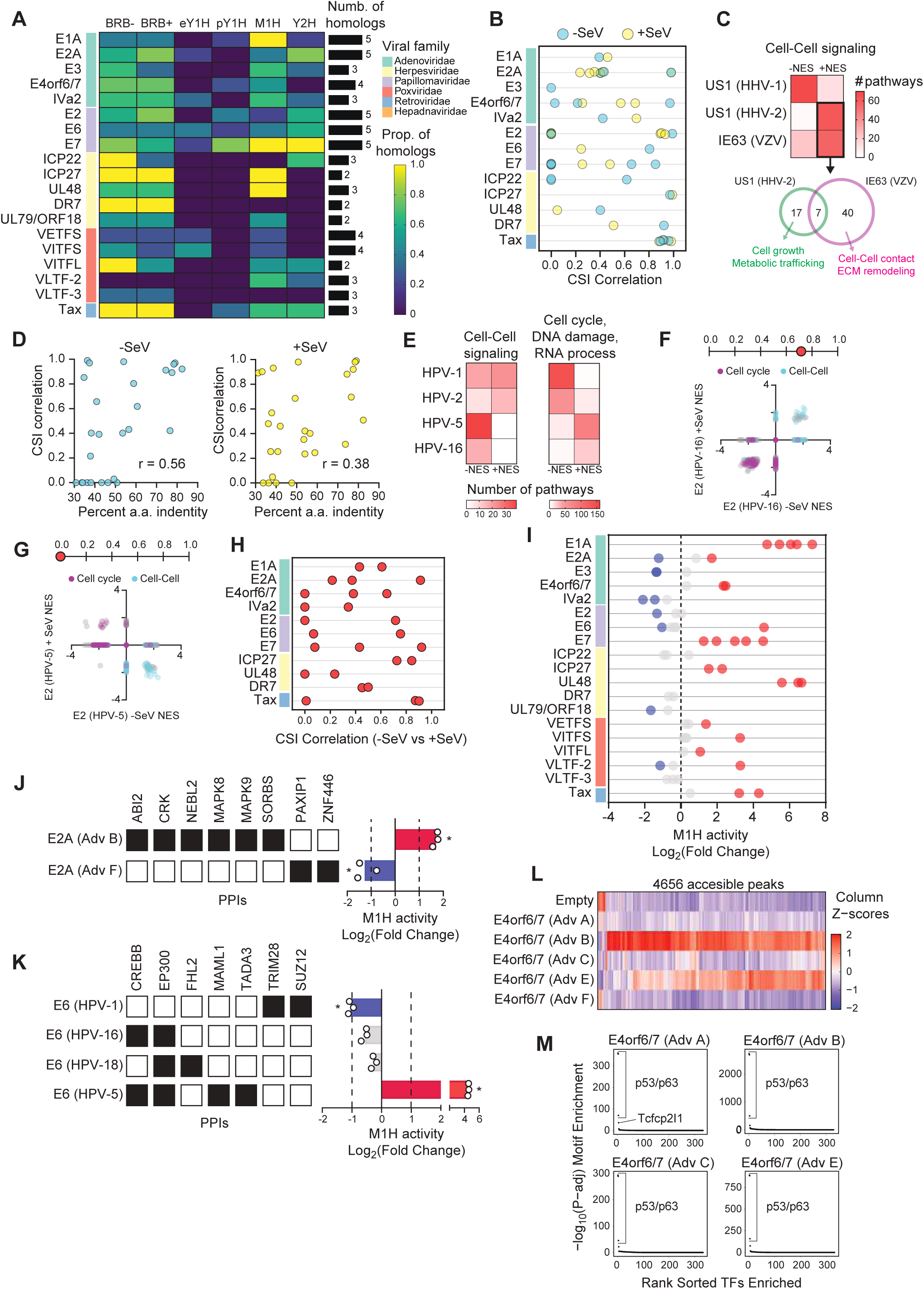
Functional variation across vTR homologs. (A) Survey of data available for sets of homologous vTRs. Cell color gradient indicates the proportion of homologs in each data set. Black bars indicate the total number of vTR homologs per set. BRB-, BRB-seq in resting cells; BRB+, BRB-seq in Sendai Virus-infected cells; eY1H, enhanced yeast one-hybrid; pY1H, paired yeast one-hybrid; M1H, mammalian one-hybrid; Y2H, yeast two-hybrid. (B) CSI correlation scores for pairs of vTR homologs. Blue and yellow dots indicate comparisons from uninfected or Sendai virus-infected cells. (C) Comparison of up- and down-regulated Cell-Cell signaling pathways between HHV ICP22 homologs. Color gradient indicates the number of pathways affected in uninfected conditions. Venn diagram represents the shared and distinct pathways upregulated by both US1 (HHV-2) and IE63 (VZV). NES, normalized enrichment score. (D) Scatter plots representing the relationship between the percent amino acid identity between homologs and CSI correlation score. Left plot, uninfected cells; Right plot, infected cells. Pearson r values are indicated in the plot. (E) Comparison of up- and down-regulated pathways between HPV E7 homologs. Color gradient indicates the number of pathways affected in uninfected conditions. (F-G) Scatter plots comparing the NES of pathways dysregulated by HPV-16 E2 (F) and HPV-5 E2 (G) in uninfected versus Sendai Virus-infected conditions. Selected pathways are colored. Number lines indicate the CSI correlation score for the given E2 homolog in uninfected versus SeV-infected cells. (H) CSI correlation scores for vTR homologs in uninfected versus SeV-infected cells. (I) Transcriptional activities of vTRs across homolog groups via M1H assays using a moderate promoter. Red and blue points indicate activating (L2FC ≥ 1) and repressing (L2FC ≤ −1) vTRs, respectively. (J-K) PPIs (left) and M1H activity (right) for two Adenovirus E2A (J) and four HPV E6 (K) and homologs. (L) Heatmap of z-score normalized chromatin accessibility at 4,656 peaks that are differentially accessible (FDR□≤□0.1 and log2FC□≥□1) via PROD-ATAC comparing E4ORF6/7 homologs from five Adenoviruses. Peak columns are hierarchically clustered. (M) Representative DNA motif enrichment within increased differentially accessible peaks for four Adv E4ORF6/7 homologs. Displayed values were calculated using hypergeometric tests and plotted relative to the ranked order of all queried motifs.

As expected, CSI transcriptional similarity is generally correlated with amino acid sequence identity between homologs (**Figure 6D**). However, there are several exceptions involving dissimilar vTR homologs affecting similar pathways. For instance, while E7 from HPV-5 and HPV-16 only share 35% amino acid identity, their CSI similarity is 0.85. Similarly, E7 from HPV-1 and HPV-2 have a CSI similarity of 0.66 with a sequence identity of 41%; however, these E7 proteins affect different pathways than those affected by E7 from HPV-5 and HPV-16, including cell-cell signaling, cell cycle, DNA damage response, and RNA processing (**Figure 6E**). We also found that the highly dissimilar E2 from HPV-16 (high-risk of cancer) and HPV-5 (low-risk) elicit a very similar transcriptional response in unstimulated cells (31% sequence identity; CSI = 0.99). However, while HPV-16 E2 maintains a robust transcriptional effect upon SeV infection, upregulating extracellular signaling and downregulating cell cycle pathways, infection flips the HPV-5 E2 transcriptional effect (**Figure 6F-G**). These differences in responses to SeV infection were also observed for other sets of vTR homologs (**Figure 6H**). Altogether, this suggests that different viral species may share functions between some but not other vTR homologs, potentially contributing to differences in infection severity and outcomes.

hTF effector domains evolve rapidly, often resulting in differences in activity levels or sign between homologs.^73^ To determine whether these differences are observed for vTRs, we compared the M1H transcriptional activity within homolog groups (**Figure 6I**). We identified several groups containing both activating and repressing vTRs such as HPV E6 and adenovirus E2A (**Figure 6I**). These homologs showed marked PPI differences (**Figures 6J-K**). The activating adenovirus B E2A interacts with proteins commonly associated with transcriptional activation, including MAPK8 and MAPK9, whereas the repressive adenovirus F E2A interacts with chromatin modulator PAXIP1 and transcription factor ZNF446, which contains a strong repressive KRAB domain (**Figure 6J**). Similarly, repressive HPV-1 E6 associates with co-repressors TRIM28 and SUZ12 (**Figure 6K**). Interestingly, E6 from both HPV-16 and HPV-18 do not exhibit strong transcriptional activity by M1H assays, despite interacting with strong coactivators such as CREBBP and EP300 (**Figure 6K**). In comparison, HPV-5 E6 is highly active and associates with CREBBP, EP300, as well as the coactivators MAML1 and TADA3 (**Figure 6K**). MAML1, typically linked to NOTCH signaling, binds CREBBP and enhances histone acetyl-transferase activity,^99^ while TADA3, a component of the PCAF complex, bridges EP300 and CREBBP.^100^ This suggests that MAML1 and TADA3 may help stabilize EP300/CREBBP function and support the transcriptional activation conferred by HPV-5 E6. Together, these data suggests that vTR homologs can have opposite transcriptional effects driven by differences in interactions with coactivators, corepressors, and hTFs.

Certain homolog groups, including adenovirus E1A and E4ORF6/7, HPV E7, and herpesvirus VP16 activate transcription, though at varying magnitudes. Yet, homologs within each of these groups affect the expression of different pathways (**Figure 6B** and **2A**), likely by regulating different target genes. Indeed, homologs of HPV E6 and E7 show different DNA binding profiles by Y1H assays (**Figure 3A**). In addition, we found marked differences in chromatin accessibility between homologs of adenovirus E4ORF6/7 by PROD-ATAC, with adenovirus B E4ORF6/7 exhibiting the broadest and strongest increase in accessibility (**Figure 6L** and **S4A**). This is also consistent with adenovirus B E4ORF6/7 affecting the expression of ∼5-fold as many genes as other E4ORF6/7 orthologs (**Figure S4B**). Despite differences in genomic position and degree of accessibility, regions of open chromatin detected upon expression of E4ORF6/7 homologs are similarly enriched for p53/p63 motifs, suggesting that binding differences may be linked to differences in PPI strengths, cooperative DNA binding, or pioneering effects (**Figure 6M**). Collectively, these results show that although vTR homologs often share certain functions, they frequently diverge in others, leading to distinct patterns of host target engagement, transcriptional outcomes, and possibly other molecular activities.

### vTRs regulate genes associated with disease risk

Viral infections have long been associated with different chronic diseases, including autoimmune and neurological disorders, through molecular mimicry, bystander activation, immune dysregulation, and alteration of immune tolerance.^101,102^ For instance, numerous studies have shown that EBV EBNA2 is recruited to the risk loci of multiple autoimmune diseases through interactions with hTFs such as RBPJ, leading to host gene dysregulation.^18,103,104,105^ To determine whether this is a unique feature of EBNA2 or whether other vTRs may also alter the expression of genes at disease risk loci, we evaluated the overlap between genes modulated by each vTR and risk loci from genome-wide association studies (GWAS). In total, we identified 40 vTRs with significant associations (FDR < 0.05) with at least one disease, corresponding to autoimmune, cardiovascular, respiratory, and neurological disorders (**Figure 7A**).

**Figure 7:**
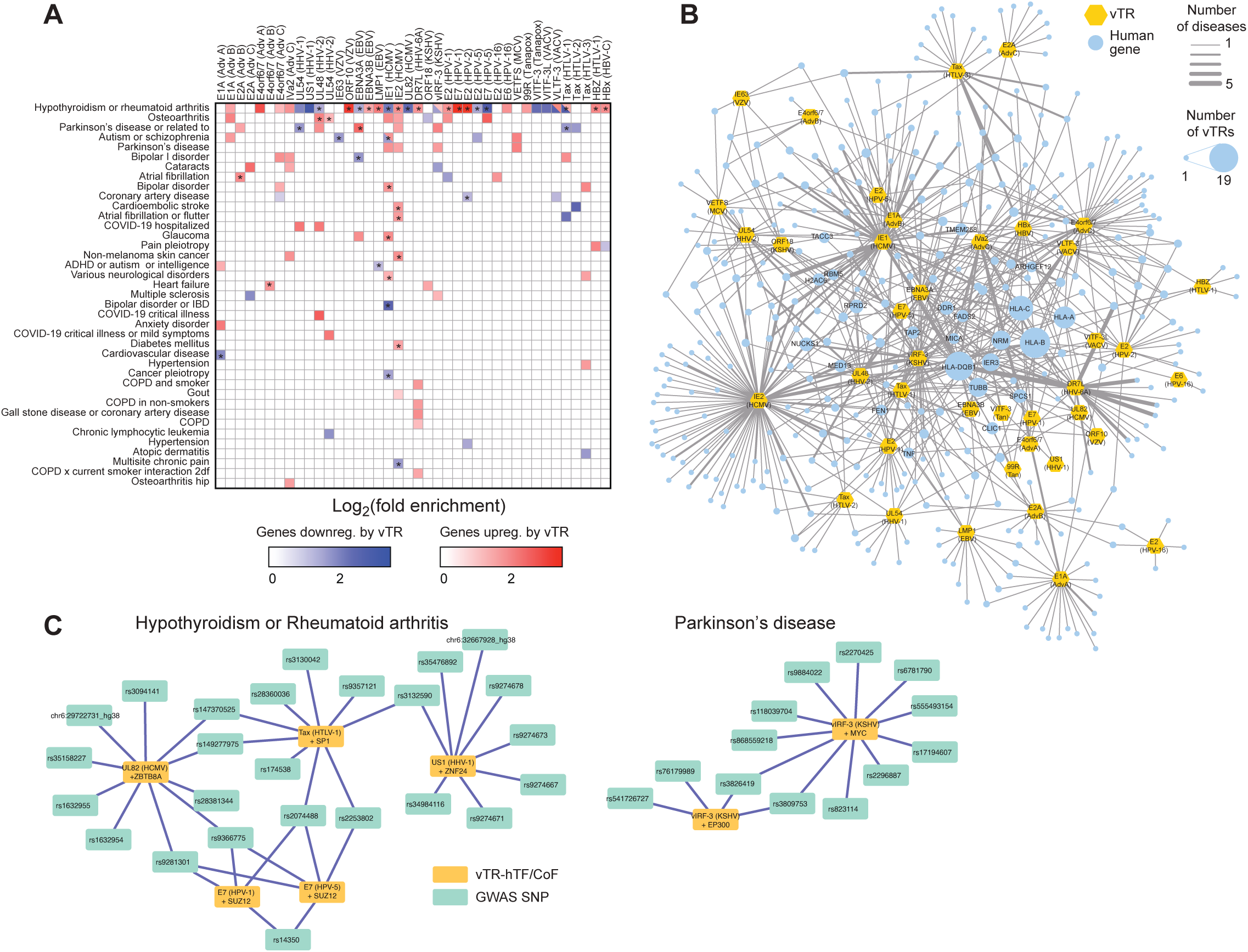
Association between vTRs and disease. (A) Fold enrichment of disease risk loci that overlap with genes up- or down-regulated by the expression of different vTRs (gene transcription start site +/- 1 kb). Only associations with an FDR < 0.05 are shown. Asterisks indicate cases where the virus expressing a particular vTR has been associated with the corresponding disease in the literature. (B) Network connecting vTRs and differentially expressed genes that are associated with disease risk loci. Edge width indicates the number of diseases in which the gene was significantly associated. Human gene node size corresponds to the number of vTRs that dysregulate the expression of the gene. Genes dysregulated by 5 or more vTRs are labeled. (C) Network showing significant vTR–hTF/CoF–GWAS locus trios for Hypothyroidism or rheumatoid arthritis, and Parkinson’s disease. Edges represent overlap between vTR DEGs and the disease risk SNP, and between ChIP-seq peaks for the hTF/CoF and the same disease risk SNP, for pairs of vTRs and hTFs/CoFs with reported PPIs.

Of the 109 significant disease-vTR associations identified, 40 are consistent with documented links between the disease and the virus encoding the vTR (**Data S7**). Several studies have shown that HCMV-infected individuals have a higher prevalence of cardiovascular diseases.^106,107^ For instance, HCMV IE2 promotes atherosclerosis by inhibiting cell death of vascular smooth muscle cells.^108,109^ We show that genes upregulated by HCMV IE2 are enriched in those associated with cardioembolic stroke and atrial fibrillation/flutter, suggesting another potential mechanism (**Figure 7A**). In addition, genes dysregulated by HCMV IE1 overlapped with those associated with multiple neurological disorders, consistent with reports of HCMV driving these diseases and the role of IE1 in perturbing neuronal development and function.^110,111^ We also found several novel associations, including between DR7L (HHV-6A) and chronic obstructive pulmonary disease, and between UL48 (HHV-2) and COVID-19 severity (**Figure 7A**). These results suggest that vTRs extensively reprogram host transcriptional networks, positioning them as potential intermediaries that connect genetic risk variants to disease-related gene dysregulation.

The disease risk genes affected generally vary across different vTRs, even those from the same virus (**Figure 7B**). However, a few disease genes are frequently targeted by multiple vTRs. Many of these genes encode immune regulators (e.g., *HLA-A/B/C/DQB1*, *MICA*, *TAP2*, *TNF*) or components of signaling and chromatin-modifying complexes (e.g., *MED13*, *MED19*, *PBRM1*, *KANSL1*, *GATAD2A*), while others are metabolic genes (e.g., *FADS1/2*). The recurrent modulation of these loci by multiple vTRs suggests that viral infection may amplify or unmask the effects of genetic variants at these sites, thereby influencing disease penetrance or severity. To determine whether the associations between vTRs and disease loci could be mediated by human regulatory factors, we integrated vTR–disease associations with hTF/cofactors ChIP-seq binding at GWAS loci for vTRs and hTFs/cofactors known to interact by PPIs. We detected 11 significant vTR–hTF/cofactor–disease trios (**Data S8**), including several involving hypothyroidism or rheumatoid arthritis, and Parkinson’s disease. Many of these trios contained overlapping SNPs between the vTR-associated loci and hTF/CoF binding sites (**Figure 7C**). For instance, vIRF-3 (KSHV) dysregulates the expression of several genes in proximity of Parkinson’s disease risk loci which also show recruitment of MYC and EP300, two proteins reported to interact with vIRF-3. Together, these results suggest that some vTR effects at disease loci may be mediated through vTR recruitment by human transcriptional regulators as has been reported for EBNA2 and RBPJ. ^18,103,104,105^

## Discussion

We systematically examined 95 vTRs across diverse human viruses, assessing their effects on transcription, DNA binding, transcriptional activity, and PPIs, providing a framework to understand how these proteins rewire host cellular programs and contribute to viral replication, persistence, and pathogenesis. For instance, we observe that vTRs across viruses consistently perturb host pathways involved in proliferation, metabolism, DNA damage, and immune response, all of which contribute to viral replication and survival. This illustrates how vTRs act as powerful modulators of host cell states, enabling viruses to simultaneously influence multiple host pathways and drive broad shifts in cellular behavior.

vTRs use different mechanisms to target these pathways. Although in some cases this is achieved by direct binding to DNA, in most cases the transcriptional effects are likely driven by hijacking endogenous regulatory proteins. One mechanism involves indirect recruitment by tethering to hTFs, consistent with findings that vTR ChIP-seq peaks are enriched for binding sites of hTFs that physically interact with the vTR.^6,18,20^ In other cases, vTRs cooperate with hTFs, which could re-direct hTFs to other genomic locations or change their activity (e.g., IRF1-E7). A less explored strategy involves vTRs preventing the binding of hTFs that regulate cellular programs that interfere with viral replication (e.g., HPV E7 antagonizing E2F4). Together, these strategies enable vTRs to rewire existing transcriptional programs that control specific biological processes, which is more parsimonious than creating entirely new programs through vTR binding recruitment to multiple loci. Interestingly, some vTRs function through multiple mechanisms (e.g., directly binding to DNA, cooperating with hTFs, and antagonizing other hTFs), which could be leveraged by viruses with compact genomes to rewire diverse pathways using few vTRs.

Many vTRs are multifunctional proteins that can affect host pathways through a variety of mechanisms, such as targeted degradation or signaling inhibition. For example, in addition to exhibiting potentially direct and cooperative DNA binding, antagonism of hTFs, and intrinsic transcriptional activity, HPV E6 promotes p53 degradation, whereas HPV E7 promotes the degradation of RB1.^112,113^ Thus, some of the transcriptional signatures we observe are likely a combination of direct transcriptional and indirect effects. Further, viruses also encode proteins that disrupt host pathways independently of transcriptional control. These include factors that promote protein degradation (e.g., KSHV K3 and K5), mimic host molecules (e.g., HCMV US28, HIV-1 Gp120), or scaffold host signaling complexes (e.g., HHV-1/2 UL46).^114–116^ Therefore, vTRs may buffer, enhance, or act orthogonally to these signaling modulators within multi-layered regulatory systems.

We found that vTRs from the same virus can have both overlapping and distinct roles in modulating host pathways. Although viruses have compact genomes with relatively few protein-coding genes, they may preserve functional redundancy for several reasons. Redundant vTRs can provide continuity in host regulation across different stages of infection (e.g., early and late vTRs both suppress immune pathways) or serve as a failsafe to counteract antiviral defenses that disable individual proteins. vTRs can also exhibit a division of labor, either by targeting different pathways that affect the various processes needed for viral replication or persistence, targeting parallel pathways that converge on the same cellular process, or by having opposing effects on the same pathway depending on context. These patterns suggest that redundancy and specialization among vTRs constitute complementary strategies that enable viruses to exert robust and fine-tuned control over their environment.

vTRs from unrelated viral species with no structural homology can produce strikingly similar effects on host pathways, underscoring widespread selective pressure to target conserved ‘pressure points’ in host biology. Conversely, homologous vTRs from closely related viruses often differ in function and mechanism. While broad regulatory agreement is shaped by the fundamental requirements for viral replication and survival, factors such as tropism can drive diversification of otherwise similar vTRs. These functional differences among homologs may also contribute to differences in pathogenicity, as seen in E2/E6/E7 across low- and high-risk HPVs or in the Tax proteins of HTLV-1, -2, and -3.^117–120^

Our analysis uncovered widespread associations between disease risk loci and genes modulated by diverse vTRs, indicating that interactions between viral activity and host genetic variation may be far more pervasive than previously appreciated. Although determining causality will require longitudinal cohorts, integrated viral-host genomic and transcriptomic datasets, and functional studies in primary human systems, the patterns we observe suggest that vTRs often modulate host regulatory networks that contribute to disease susceptibility. In this framework, viral infection may act as a broad environmental trigger that functionally engages susceptibility loci, influencing disease penetrance, severity, and heterogeneity. Given the high prevalence of many of these viruses, these findings underscore the importance of systematically incorporating viral serostatus into GWAS designs to more accurately assess the combined effects of inherited and viral factors on complex disease risk.

Altogether, our findings highlight vTRs as versatile regulators that leverage diverse molecular mechanisms yet consistently converge on core host pathways essential for replication, persistence, and immune evasion. Beyond defining these principles, this work establishes a broadly applicable resource for the virology community. The multi-modal datasets generated will enable researchers studying a specific vTR or virus to contextualize their findings within a comparative framework. Such cross-virus comparisons can reveal whether vTRs not tested in our study, a newly identified vTR or a vTR ortholog from specific strains act through strategies already observed in our datasets. This may suggest candidate host proteins, pathways, mechanisms of DNA recruitment, or links to disease to prioritize for in-depth characterization. Likewise, our multimodal profiles can help reinterpret viral infection phenotypes by pointing to underlying vTR drivers. By mapping how vTRs engage host gene regulatory networks, this work identifies conserved regulatory nodes that may offer new therapeutic targets in both acute and chronic infections. Together, our work creates a foundation for future investigations into the roles of vTRs in viral infection and disease.

## Resource availability

### Lead contact

Requests for information should be directed to Juan I. Fuxman Bass.

### Materials availability

All unique/stable reagents generated in this study will be made available from the lead contact with a completed material transfer agreement.

### Data and code availability

Original code is available at https://github.com/FuxmanBass-lab/vTRs. Any additional information required to reanalyze the data reported in this paper is available from the lead contact upon request.

## Acknowledgments

The work was supported by NIH grants R35GM128625 (to J.I.F.B.), R35GM133658 (to S.S.Y.), R35GM137836 (to N.S.), U01CA232161 (to J.I.F.B., M.V., and M.L.B.), R01HG010730, U24HG013078, R01NS099068, and R01AI024717 (to M.T.W.), and Alfred P. Sloan Scholar Research Fellowship grant FG-2018-10723 (to N.S.). M.Y.E and P.T. were supported by NIH training grant T32GM150533. P.D. was supported by NIH training grant T32AR069512. S.K. was supported by NIH grant K99HG013675. S.R. was supported by the Little Warrior Foundation and the Vilas Associate Fellowship.

## Author contributions

Conceptualization, J.T.R., X.L., J.I.F.B.; validation, J.T.R., X.L., L.F.S-U., B.E.; formal analysis, J.T.R., X.L., L.S-U., M.Y.E., J.E.C., P.D., B.E., G.M-E., Z.L., P.T., A.B., S.S., T.H., J.I.F.B.; investigation, J.T.R., X.L., A.B., C.S., J.E.C., R.L., B.E., Y.L., K.S-F., S.K., S.S., N.S.., C.L., L.M-C., J.I.F.B.; resources, J.I.F.B., M.V., S.R., M.L.B., M.T.W.,; data curation, J.T.R., X.L., J.I.F.B., M.W.; writing – original draft, J.T.R., J.I.F.B.; writing – review and editing, J.T.R., J.I.F.B., N.S., S.Y., S.R., M.T.W., A.B., L.F.S-U., M.L.B., S.K., M.V.; visualization, J.T.R., M.Y.E., B.E., A.B., J.E.C., J.I.F.B.; supervision, J.I.F.B., S.R., M.V., M.C., M.L.B., M.T.W.; project administration, J.I.F.B.; and funding acquisition, J.I.F.B, M.V., S.R., M.L.B., M.T.W.

## Declaration of interests

The authors declare no competing interests.

## Supplementary Figures

**Figure S1:**
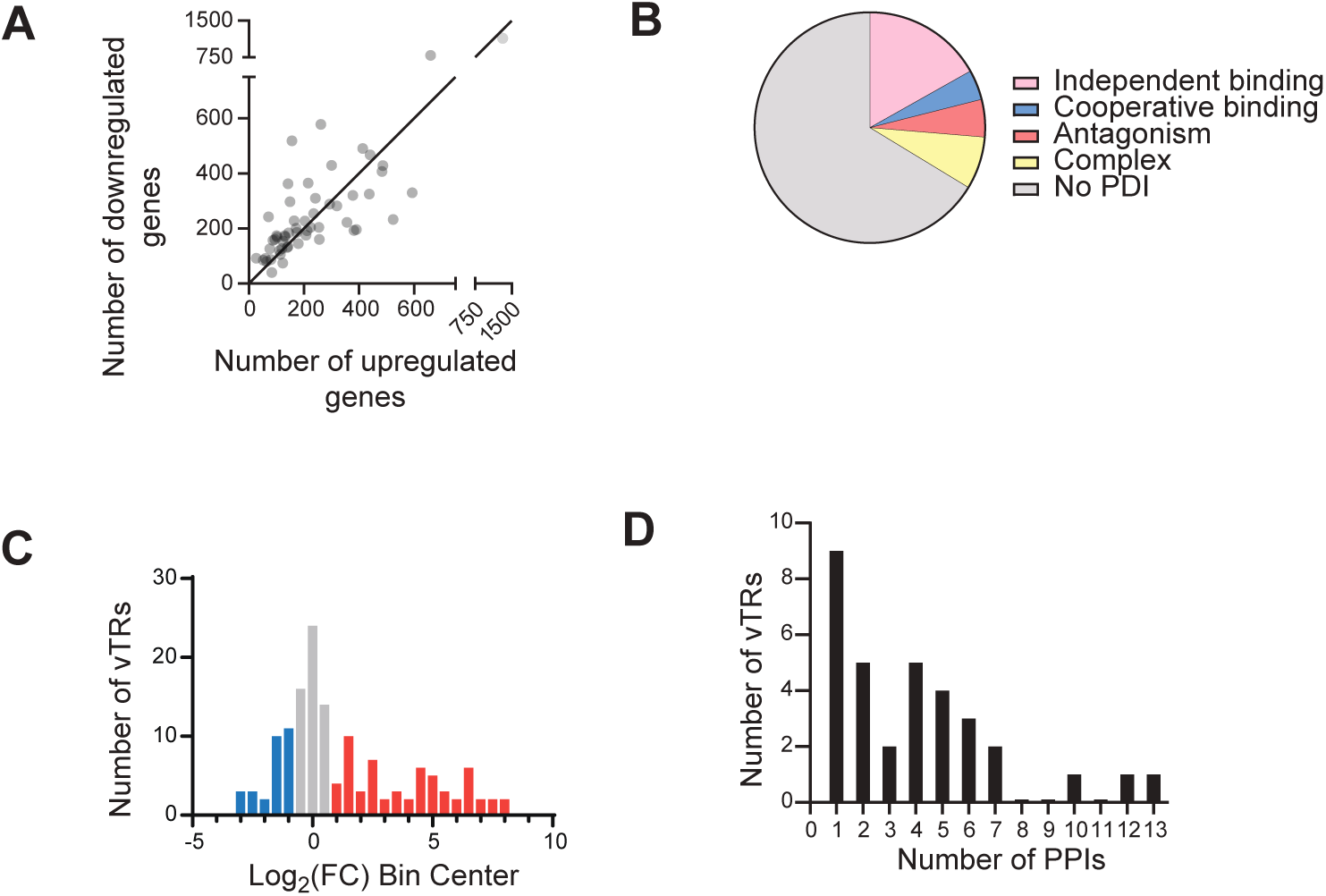
Overview of datasets generated using vTR1.0. (A) Number of differentially expressed genes for each vTR successfully tested via BRB-seq. Each point represents the number of up- and down-regulated genes driven by the expression of each vTR (|Log2 FC| ≥ 0.5 and p ≤ 0.05). (B) Proportion of protein-DNA interactions (PDIs) determined for vTRs tested using eY1H and pY1H assays that were classified as independent (does not require a partner), cooperative (binding is enhanced by a partner), antagonistic (one protein prevents the binding of the other), or complex (binding modality depends on the target sequence). (C) Histogram of distribution of transcriptional activity for vTRs tested using M1H assays, relative to vector control. Activators (Log2 FC ≥ 1) are colored red and repressors (Log2 FC ≤ - 1) are colored blue. (D) Distribution of protein-protein interactions (PPIs) detected between vTRs and human proteins using Y2H assays.

**Figure S2:**
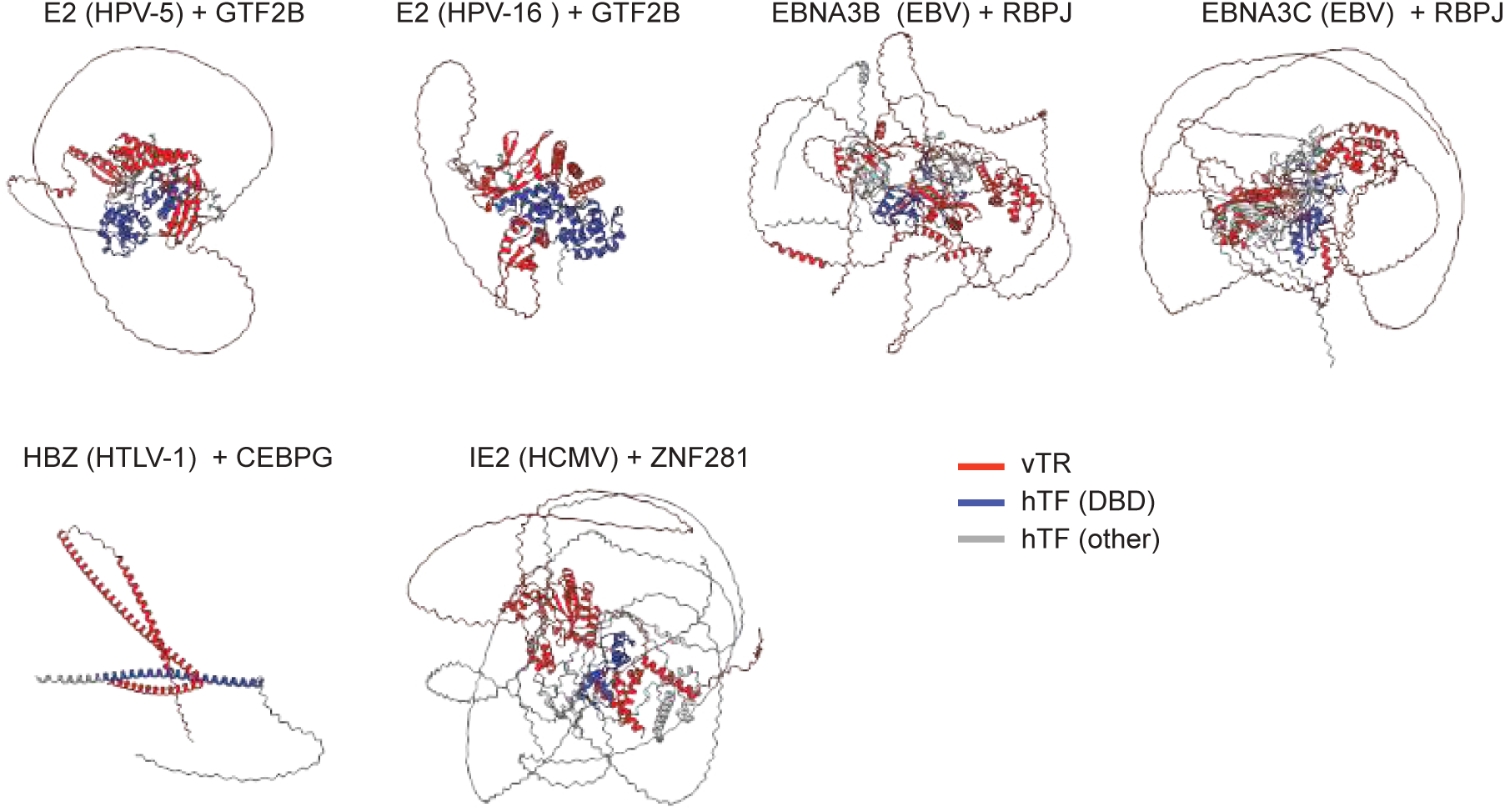
Alphafold3 models between vTRs and hTFs. Alphafold3 models of vTRs-hTF pairs where the vTR antagonizes the binding of a hTF to DNA. vTR is shown in red, the hTF DBD is shown in blue, and non-DNA regions are shown in gray.

**Figure S3:**
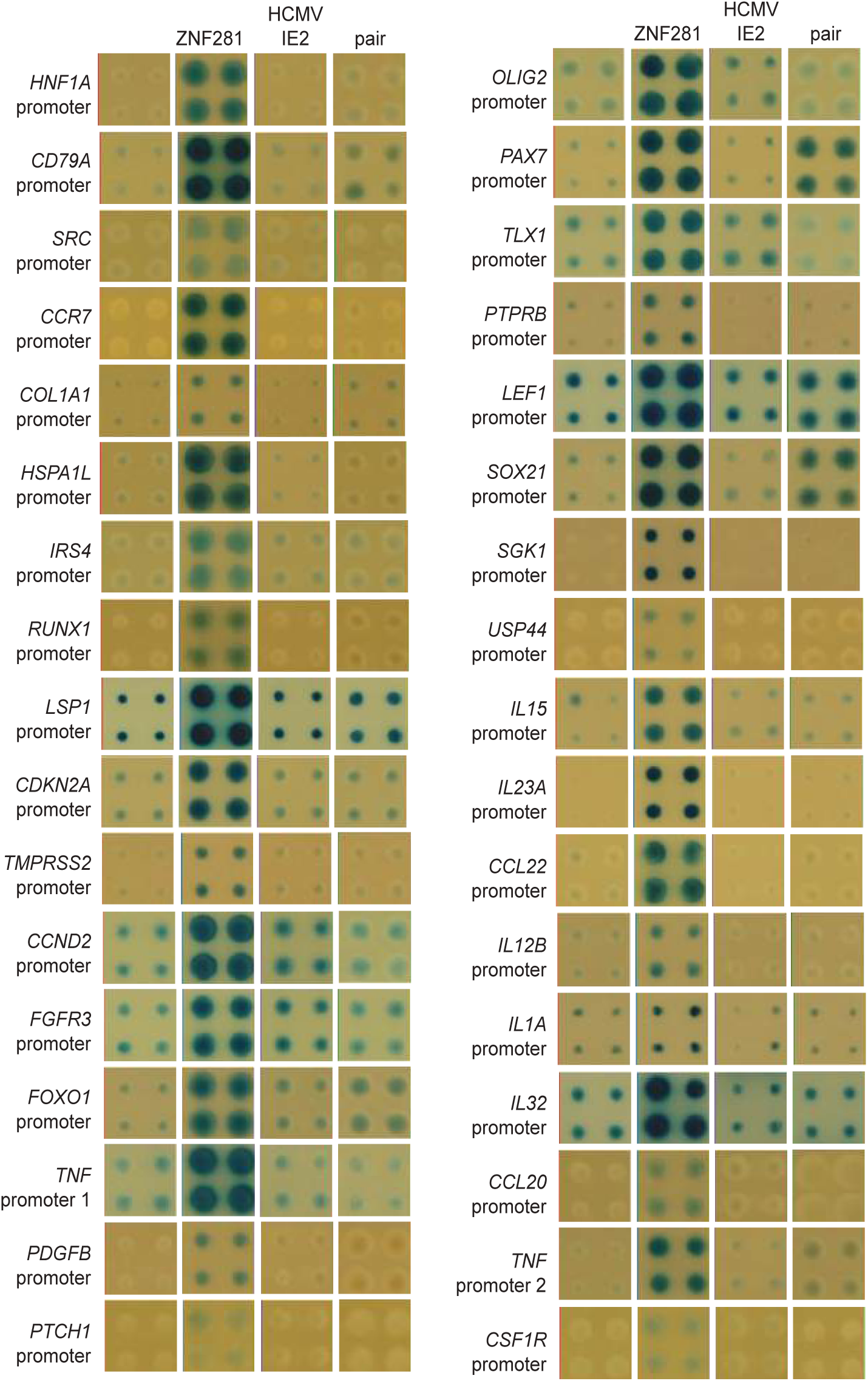
pY1H antagonistic interactions between ZNF281 and IE2 (HCMV) Antagonistic pY1H interactions involving IE2 (HCMV) and hTF ZNF281. Italicized gene names indicate the promoter tested. Yeast strains shown: empty vector control, ZNF281 only, IE2 only, and ZNF281-IE2 pair.

**Figure S4:**
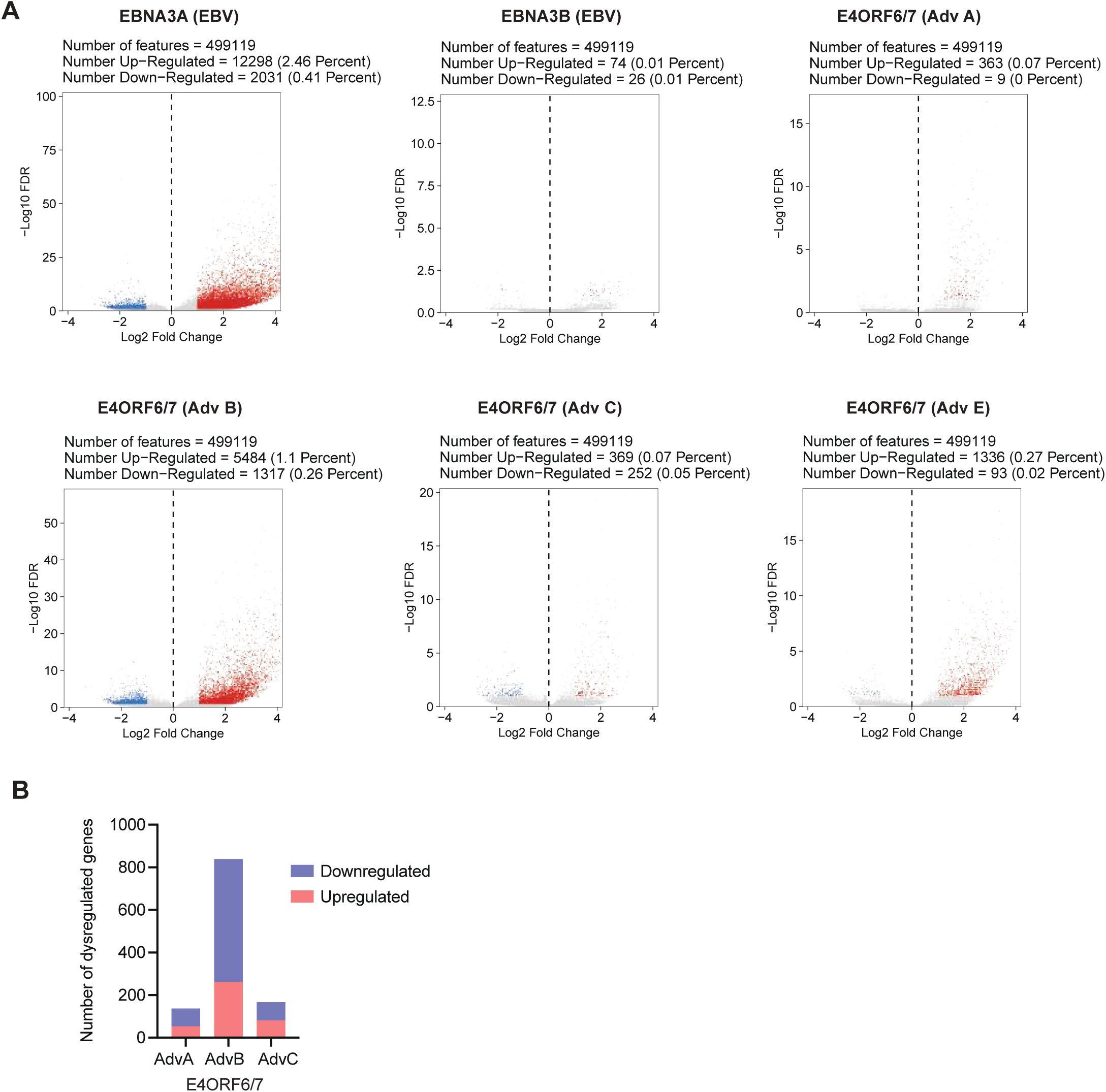
Differentially accessible chromatin regions identified by PROD-ATAC. (A) Volcano plots depicting the −log(FDR) versus log2FC for each called peak comparing pseudobulk replicates of the corresponding vTR to the Empty Vector control. Peaks with increased accessibility (FDR□≤□0.1 and log2FC□≥□1) are red, and peaks with decreased accessibility (FDR□≤□0.1 and log2FC□≤□−1) are blue. (B) Number of differentially expressed genes driven by expression of E4ORF6/7 from different adenoviruses.

## Methods

### EXPERIMENTAL APPROACHES

#### Generation of vTR1.0 entry clone collection

95 vTR ORF sequences (**Data S1**) were synthesized as gene fragments (Twist Bioscience) and subsequently cloned into pDONR221 by Gateway BP reaction (Thermo Fisher Scientific) followed by transformation into *E. coli* DH5□. Transformed *E. coli* were then plated onto LB agar containing 50 ug/mL Kanamycin overnight at 37 C. Colonies for each transformation were confirmed by Sanger sequencing and expanded to generate our vTR entry clone collection.

#### BRB-seq experiments

To determine the transcriptomic effects of vTRs in human cells, we performed bulk RNA barcoding and sequencing (BRB-seq) in HEK293T cells expressing individual vTRs. Expression vTR clones were generated via Gateway LR cloning into the pEZY3 constitutive expression vector (Addgene plasmid #18672). Following transformation, plasmids were extracted, and insertion was confirmed via PCR and gel electrophoresis. The full list of vTRs tested by BRB-seq is provided in **Data S2**.

For BRB-seq experiments, HEK293T cells were cultured in DMEM media supplemented with 10% FBS and seeded in two 96-well plates at a density of 18,000 cells/well. The following day, cells were transfected in quadruplicate using Lipofectamine 3000 (Thermo Fisher Scientific) with 80 ng of the pEZY3-vTR plasmids, as well as empty pEZY3 vector controls. After 24 hours, cells were either infected with 4.0 × 10□ CEID₅₀/mL Sendai virus (VR-907, ATCC), or left uninfected. RNAs were harvested 2 days post-transfection, a time point that we showed has minimal interferon pathway activation due to transfection. Sequencing libraries were prepared using the MERCURIUS DRUG-seq kit (Alithea Genomics) according to manufacturer’s instructions and as described previously.^24^ Briefly, HEK293T cells were permeabilized using a cell lysis buffer on ice 15 min with periodic agitation, followed by centrifugation to clear cellular debris. Individual RNA-containing lysates were then reverse-transcribed using barcoded oligo-dT primers such that the resulting cDNA from each individual sample was uniquely barcode-labeled. cDNA from all samples in a single plate were then pooled and purified using the column-based Zymo Clean & Concentration Kit according to manufacturer’s instructions (Zymo, D4014). The pooled cDNA was then cleared of non-incorporated primers via exonuclease digestion, followed by second strand synthesis and quantification of ds-cDNA via Qubit HS-DNA kit (Thermo Fisher). Tagmentation was then performed on 20 ng cDNA, followed by library indexing and amplification using standard Illumina NGS adapter sequences. Indexed libraries were sequenced at a depth of 5 million raw reads per sample on an Illumina Novaseq platform.

#### Enhanced yeast one-hybrid (eY1H) assay

To detect PDIs between vTRs and cytokine gene promoters, eY1H assays were performed as previously described.^121–123^ vTRs prey clones were generated via Gateway LR into pDEST-AD2µ (Walhout lab) such that each vTR was fused with the yeast Gal4 Activation Domain. vTR-prey strains were generated via transformation into Y□1867 yeast as previously described.^121,124^ Yeast were inoculated in 1 L liquid YAPD media at a concentration of OD600 = 0.15 and incubated at 30°C shaking at 200 rpm until they reached OD600 = 0.4-0.6, washed with sterile water, and washed again with TE + 0.1 M lithium acetate (TE/LiAc). Yeast were resuspended in TE/LiAc with salmon sperm DNA (ThermoFisher #15632011) at a dilution of 1:10 before adding 250 ng of the vTR clone. Six volumes of TE/LiAc + 40% polyethylene glycol were then added, and samples were mixed gently ten times. Yeast were then incubated at 30°C without shaking for 30 minutes followed by 42°C for 20 min, then resuspended in sterile water. Transformed yeast were plated in selective media lacking tryptophan to select for transformants. vTR prey yeast strains were confirmed by Sanger sequencing.

DNA-bait strains corresponding to 112 human cytokine promoters were previously generated using the Y1HaS2 yeast strain.^25^ Bait-strains carry two integrated copies of the promoter element upstream of HIS3, which allows yeast to grow in the absence of histidine and overcome growth inhibition by 3-amino-1, 2, 4-triazole (3AT), and LacZ which causes yeast colonies to turn blue in the presence of 5-bromo-4-chloro-3-indolyl-beta-D-galacto-pyranoside (X-gal).

eY1H was performed as previously described ^121–123^ using a high-density array ROTOR robotic platform (Singer Instruments). The vTR yeast array and cytokine promoter yeast strains were mated pairwise and spotted on permissive agar media for 1 day at 30°C followed by selection on agar plates lacking Uracil and Tryptophan for 2 days to select diploid yeast. Selected diploid yeast were then transferred to selective media agar plates lacking uracil, tryptophan, and histidine, with 5mM 3AT and 320 mg/L X-gal. These readout plates were imaged at 2-, 3-, 4-, and 7-days post-plating. Interactions were tested in quadruplicate and were manually curated by 2 researchers. Only vTR-promoter pairs that resulted in growth and blue color for at least two of the four colonies were considered positive (**Data S3**). At least 90% of interactions were detected with all four colonies.

#### Paired Yeast one-hybrid (pY1H) assay

To detect cooperative binding and antagonism between vTRs and hTFs, pY1H assays were performed as previously described.^26^ Prey strains expressing 113 vTR-hTF pairs were generated using a Y□1867 strain where the LEU2 gene had been disrupted (Y□1867Δleu2) to accommodate transformation with plasmids containing both TRP1 and LEU2 selection markers.^26^ This array vTR-hTF array also included clones with single vTRs or hTFs expressed. DNA-bait strains corresponding to 39 human cytokine promoters and 83 cancer-related gene promoters were previously generated using the Y1HaS2 yeast strain.^25,125^

pY1H was performed using a high-density array ROTOR robotic platform (Singer Instruments) as previously described. ^26^ Briefly, the vTR-hTF yeast array and human promoter yeast strains were mated pairwise (in quadruplicate) on permissive medium agar plates and incubated at 30°C for 1 d. Mated yeast were then transferred to selective medium agar plates lacking uracil, leucine, and tryptophan to select for successfully mated yeast and incubated at 30°C for 2 d. These selection plates were imaged and analyzed to identify array locations with failed yeast growth, which were then removed from further analysis. Diploid yeast were finally transferred to selective medium agar plates lacking uracil, leucine, tryptophan, and histidine, with 5 mM 3AT and 320 mg/liter X-gal. These Readout plates were imaged 2, 3, 4, and 7 d after final plating.

Yeast plate images were processed and visualized using DISHA (Detection of Interactions Software for High-throughput Analyses) software.^26^ vTR-hTF pair strains were sorted based on each index (cooperativity, antagonism index 1, and antagonism index 2) separately. Images were then manually analyzed to identify cooperative and antagonistic interactions. To call an interaction, the following criteria was used: 1) vTR-hTF pair, vTR, and hTF yeast strains all showed growth in the mating selection plates before transfer to readout plates; and 2) on readout plates, ≥3 out of 4 replicate colonies were uniform for vTR-hTF pair, vTR, and hTF yeast strains. For cooperative interactions, vTR-hTF pair yeast showed a strong or moderate reporter activity relative to the empty-empty strain, and vTR and hTF yeast showed no or only weak reporter activity. For antagonistic interactions, vTR and/or hTF yeast showed a strong or moderate reporter activity relative to the empty-empty strain and the vTR-hTF pair yeast showed no or only weak reporter activity. Cooperative and antagonistic interactions are provided in **Data S4**.

#### Mammalian one-hybrid (M1H) assays

To measure vTR transcriptional activity, M1H assays were performed in HEK293T cells.^75,125^ In this assay, vTR fused to the Gal4 DNA-binding domain are recruited to the upstream activating sequence (UAS) upstream of the firefly luciferase gene. Changes in expression of firefly luciferase are reflective of the endogenous activating or repressing activity of the vTR.

vTRs were cloned into the DB-pEZY3 vector via Gateway LR such that the vTRs were N-terminally fused to Gal4 DNA-binding domain.126 M1H assays were performed using pGL4.23 luciferase reporter constructs. For low-background M1H assays, a 4xUAS + minimal promoter construct was used, as described. ^75,125^ We also performed M1H assays using a moderate background activity construct containing 4xUAS + moderate promoter containing the Syn2B10 enhancer sequence previously described^126^ to detect transcriptional repression with higher sensitivity, as well as activation.

HEK293T cells were cultured in DMEM media supplemented with 10% FBS and plated in 96-well white opaque plates (Corning) at a density of 10,000 cells/well and incubated at 37°C, 5% CO2 for one day. Cells were then transfected with 80 ng of vTR (DB-pEZY3) plasmid, 20 ng 4xUAS construct, and 10 ng of renilla luciferase plasmid as a transfection normalization control. An empty DB-pEZY3 plasmid and 4xUAS construct plasmid were co-transfected to serve as a negative control. Cells were incubated for 2 days after transfection at 37C with 5% CO2, followed by measurement of both firefly and renilla luciferase activity using the Dual-Glo Luciferase Assay System (Promega). Non-transfected cells were used to subtract background for firefly/renilla luciferase activities. Firefly luciferase activity was then normalized with renilla luciferase activity in each well. vTR activities using both reporter systems are provided in **Data S5**.

#### PROD-ATAC Pooled Library Generation

The pooled library of HEK293T cells used to generate PROD-ATAC (Protein-Coding single-cell Assay for Transposase Accessible Chromatin) chromatin accessibility profiles were produced for the described set of vTR sequences as previously described in detail (Frenkel et al, 2024). Briefly, 3-wells of a 6-well plate were seeded with 400,000 HEK293T-B4 attP-containing landing pad cells and left to attach overnight in DMEM media supplemented with 10% FBS. At approximately ∼50% confluency, each well was transfected (using Lipofectamine 3000 in Opti-MEM) with 1,500 ng donor attB library, encoding each of the relevant vTR sequences, and 100 ng of pCAG-NLS-HA-Bxb1 (Addgene, 51271). Liposome-containing media was aspirated 18 hours post-transfection and replaced with fresh complete media containing 0.8 ug/ml puromycin. After 3 days of puromycin selection, the 3-wells were pooled into a single T-25 flask. Passages were subsequently performed every 2 days (when cells reached ∼70% confluency). At 12 days post-selection start, several cell aliquots were frozen (5% DMSO v/v) and stored in the vapor phase of a liquid nitrogen dewar.

#### PROD-ATAC Screening

In preparation for the PROD-ATAC sequencing run, the HEK293T cell library encoding the described vTR sequences was seeded and allowed to attach overnight in DMEM media supplemented with 10% FBS. Cells were treated with 2 ug/ml doxycycline to induce library expression for 96 hours, with a single split performed at 48 hours post-induction. After this point, the PROD-ATAC single cell sequencing library generation procedure was performed as previously described using 10x Genomics scATAC kits with v2 Chemistry, with solely two deviations: 15,000 nuclei (rather than 10,000) were targeted for capture per 10x chip lane, and 8 total lanes were used in the assay.^127^

#### Yeast two-hybrid (Y2H) assays

Y2H assays were performed as previously described.^75,128^ vTRs were cloned into pDESTDB-X vectors via Gateway LR such that each vTR was fused to the yeast Gal4 DNA-binding domain (DB). The competent yeast strain Y8930 (MAT□) was transformed with individual DB-vTR constructs and plated on yeast synthetic media lacking leucine to select for DB-vTR plasmid-containing yeast. Haploid DB-vTR yeast strains were tested for auto-activation of the GAL1::HIS3 reporter gene. Individual DB-vTR yeast strains were spotted on SC-Leu-His+1mM 3AT media and any strains showing growth were considered auto-activators (AAs) and removed from the collection of strains to be screened.

A primary Y2H screen was performed in which all non-AA vTR strains were tested against the hORFeome v9.1 collection of 17,472 ORF clones^128^ tagged with Gal4 activation domain (AD). DB-vTR strains were mated individually against 99 pools of ∼188 AD-ORF strains (preys). To perform the mating, fresh overnight cultures of DB-vTR strains were mixed with AD-ORF strain pools and grown overnight at 30°C in liquid rich media (YEPD). After overnight growth, the mated yeast cells were transferred into liquid SC-Leu-Trp media to select for diploids and again grown overnight at 30°C. Finally, the yeast cells were spotted onto SC-Leu-Trp-His+1mM 3AT solid media to select for activation of the GAL1::HIS3 reporter gene. In parallel, diploid yeast cells were transferred onto SC-Leu-His+1mM 3AT solid media supplemented with 1 mg/l cycloheximide to test for spontaneous DB-ORF auto-activators. After 72 hours incubation at 30°C, yeast that grew on SC-Leu-Trp-His+1mM 3AT but not SC-Leu-His+1mM 3AT+ 1 mg/L cycloheximide media were picked into SC-Leu-Trp grown overnight then processes to determine identity of interaction partners by Sanger sequencing. In total, 674 PPIs between vTRs and human proteins were identified, involving 41 vTRs.

Following this first-pass screening, vTRs were tested pairwise for interactions with the candidate human ORFs identified in the first-pass screening in a matrix format (the 41 vTRs against the union of all ORF interactors). Yeast strains corresponding to the identified human interaction partners were picked from archival glycerol stocks, cultured in liquid medium and mated one-by-one against all 41 vTRs, processed as described above and then scored. Briefly, interactors were inoculated in 200 μL of selective media and mated overnight at 30°C in 150 μL liquid rich media (YEPD). The following day, mated yeast cells were transferred to 150 μL liquid SC-Leu-Trp media to select for diploids. After overnight incubation at 30°C, 5 μL diploid yeast cells were spotted onto SC-Leu-Trp-His+1mM 3AT solid media to select for activation of the GAL1::HIS3 reporter gene as well as on SC-Leu-Trp to control for successful mating. AA tests were included by mating each Y8930:DB-ORF against a Y8800:AD-null (containing no ORF), which was included on each test plate for each individual Y8930:DB-ORF. Spots were scored for growth with scores of 0, 1, 2, 3, 4, and NA for cases where the spotting had failed. Growth scores of 0 and 1 were considered not growing. If a spot corresponding to either the pair or the corresponding AA-test did not grow on SC-Leu-Trp, the pair was scored NA. If the AA-test had a growth score of 4, the corresponding pair was scored NA. Interactions were scored positive if they had higher growth scores on SC-Leu-Trp-His+1mM 3AT solid media compared to the auto-activator test and had a growth score of at least 2; otherwise they were scored negative. All positive scored colonies were picked, lysed and the identity of the two interacting proteins was confirmed by sequencing. 86 PPIs were validated, involved in 28 vTRs. Next, we curated PPIs between non-AA vTRs and human proteins previously reported in Uniprot. All human proteins that interacted with at least one vTR, with available clones in the ORFeome, were pairwise tested against the non-AA vTRs. This led to 46 interactions, which were combined with the previous 86 PPIs for a total of 132 interactions between 33 vTRs and 67 human proteins (**Data S6**).

#### Immunoprecipitation and immunoblotting analyses

HEK293T cells stably expressing the corresponding vTRs were lysed with RIPA lysis buffer (20mM Tris-HCl, 37mM NaCl2, 2mM EDTA, 1% Triton-X, 10% glycerol, 0.1% SDS and 0.5% sodium deoxycholate) with protease and phosphatase inhibitors (Roche). Protein samples were quantified (Pierce BCA Protein Assay Kit, Thermo Fisher Scientific) and the same amount of cell lysates from vTR overexpressing stable cells and GFP vector control cells were incubated with an anti-Flag M2 Affinity Gel (Sigma) overnight at 4°C. The beads were washed 3x with TBST buffer and then boiled in sample buffer. The supernatants were subjected to SDS polyacrylamide gel electrophoresis and transferred to polyvinylidene difluoride membranes. Membranes were incubated with primary antibodies, including anti-AES (1:1,000; Proteintech) and anti-Flag (1:2,000; Sigma) and subsequent secondary antibody: horseradish peroxidase–conjugated Affinipure Goat Anti-Rabbit IgG(H+L) (1:3,000; Proteintech, catalog no. SA00001-2).

### QUANTIFICATION AND STATISTICAL ANALYSES

#### BRB-seq sequence alignment

BRB-seq reads were aligned to a custom human reference based on GRCh38 (primary assembly) with Gencode v46 annotations,^129^ augmented by a concatenated FASTA of all VTR sequences and a merged GTF that combined genome and VTR features. Genome indices were built with STAR v2.7.9a^130^ using splice junctions (–-sjdbGTFfile Gencode+VTR, –-sjdbOverhang 79). Alignment and counting were performed with STARsolo in CB/UMI simple mode to recover per-barcode gene counts (–-soloType CB_UMI_Simple; –-soloFeatures Gene; –-soloStrand Forward; –-clipAdapterType CellRanger4; –-outSAMtype BAM SortedByCoordinate; –-outSAMunmapped Within; –-outReadsUnmapped Fastx). Barcodes and UMIs were parsed with a 14-nt cell barcode and 14-nt UMI (–-soloCBstart 1 –-soloCBlen 14; –-soloUMIstart 15 –-soloUMIlen 14), matched to a provided whitelist allowing one mismatch (–-soloCBmatchWLtype 1MM), and deduplicated with directional one-mismatch UMI collapsing (–-soloUMIdedup 1MM_Directional). STAR output included comprehensive SAM attributes (NH HI AS nM CR CY UR UY CB UB GX GN sS sQ sM).

#### Pre-processing, Differential Expression and Pathways Analysis

Raw STARsolo matrices were imported into Seurat^131^ for each plate-condition, and well-level metadata were joined from barcode annotation tables. For each well, we quantified the proportion of reads mapping to the expected vTR and compared it with the most abundant vTR signal within the full vTR panel. Wells with at least 85% of reads assigned to a single vTR were flagged as highly specific and used to assess concordance with the expected assignment. For downstream analyses, we applied the following filters: (i) vTR wells were required to have at least 85% of their vTR-panel reads assigned to the expected construct and a minimum of 500,000 total mapped reads; (ii) control and vector wells were required to show no dominant vTR signal (top VTR <0.01% of total reads) and at least 500,000 total mapped reads. After filtering, counts were merged across wells and joined to Gencode v46 annotations.^129^ We retained protein-coding, T-cell receptor, and immunoglobulin genes, while excluding mitochondrial and ribosomal genes as well as lowly expressed genes. Normalization and differential expression were performed using DESeq2,^132^ with the vector set as the reference. A pooled matrix design that included batch as a covariate was chosen, as it showed high correlation between replicates while effectively reducing noise and confounding effects. Results were extracted using Benjamini–Hochberg correction (FDR < 0.1) without independent filtering. Gene Set Enrichment Analysis (GSEA) ^133^ used msigdbr to retrieve MSigDB C2:CP-REACTOME sets^134^ and was run with 10,000 permutations, Benjamini-Hochberg correction, eps=0, minGSSize=15, maxGSSize=500, and p-value cutoff reporting at 1. Pathways with a p-value > 0.05 were considered not significant.

#### Reactome Pathway Categorization

Individual reactome pathways from BRB-seq experiments were designated under any number of the following categories: cell-cell signaling, signal transduction, immune, metabolism, stress, DNA damage repair, cell cycle, gene regulation/RNA biogenesis. Pathways were categorized based on reactome pathway descriptions (https://reactome.org/) and genes present in the pathway.

#### CSI correlation calculation

To quantify sample-sample similarity at the pathway level, we constructed a matrix of normalized enrichment scores (NES) from GSEA Reactome results, setting values to zero when the corresponding pathway had a non-significant enrichment (p > 0.05). Pathways represented in fewer than three samples were excluded, and missing values were imputed as zero. Pairwise sample correlations were then computed using Pearson’s correlation across the filtered NES matrix, and the resulting correlation distances were used as input for Connection Specificity Index (CSI).^31,32^ For each sample pair (A, B), the CSI was defined as the proportion of third samples (X) for which the maximum of the distances (A–X, B–X) was smaller than the distance between A and B. This procedure was parallelized across all sample pairs, producing a symmetric CSI matrix, which was normalized by the number of possible third samples (n − 2) to yield values between 0 (no concordance) and 1 (high concordance). The resulting CSI matrix was then used for downstream clustering and comparative analyses.

#### Alphafold3 of vTR structures

Predicted three-dimensional structures of proteins were generated using AlphaFold3.^135^ Structural predictions were run through the publicly accessible AlphaFold3 interface using default parameters. Briefly, amino-acid sequences of each protein of interest were retrieved from UniProt and supplied as input to the AlphaFold3 prediction pipeline. Five structural models were produced per sequence. Predicted Local Distance Difference Test (pLDDT) scores and Predicted Aligned Error (PAE) heatmaps were used to assess overall model quality and to identify well-resolved regions suitable for interpretation. The highest-confidence monomer model was selected for downstream analysis and visualization, which was performed using UCSF ChimeraX.^136^

Protein–protein complex (heterodimer) structures were predicted using AlphaFold3’s multimer mode. Human transcription factors or TLE5 were paired with vTRs and supplied jointly as separate chains in a single multimer prediction run. Five multimer models were generated for each input pair and ranked using pLDDT, inter-chain PAE, and AlphaFold3’s interface confidence metrics (iPTM and chain-pair interface scores). The top-ranked multimer model was selected for downstream interpretation, and all structural visualization and interaction inspection were performed using UCSF ChimeraX.^136^

#### Transcriptomics comparison between SeV-stimulated and unstimulated conditions

Differential expression was performed by comparing (i) each SeV-infected vTR against the SeV-infected vector control, and (ii) SeV-infected vector controls against uninfected vector controls. Count matrices were constructed from STARsolo outputs and joined with matched metadata to ensure consistent gene and sample annotation. DESeq2 (v1.46) ^132^ was used with poscounts normalization to account for sparsity in UMI data, and significance was assessed with Benjamini–Hochberg correction.

Pathway enrichment was carried out with fgsea (v1.32)(https://doi.org/10.1101/060012) using curated Reactome sets for both comparisons. Genes were ranked by log2 fold change, restricted to those present in Reactome sets, and tested with 10,000 permutations. Non-significant pathways (FDR ≥ 0.05) were set to zero before aggregation into a summary NES matrix. To highlight construct-specific effects beyond stimulation alone, discordant pathways were defined as those enriched in opposite directions between stimulated vector controls and specific stimulated vTRs. These pathways were summarized across vTRs, and clustering of normalized enrichment scores was used to visualize the relationship between constructs and pathway-level responses.

#### RELI analysis

To identify which differentially expressed genes (DEGs) resulting from vTR expression were associated with disease risk loci, we utilized the Regulatory Element Locus Intersection (RELI) approach as described by Harley et al.^18^ RELI statistically tests for enrichment of GWAS risk loci in user-supplied genomic regions or gene lists, through permutation and comparison to matched controls. We input the list of up- or down-regulated genes resulting from the expression of each vTR into RELI and assessed the enrichment of risk-associated loci from published GWAS datasets proximal to these DEGs (transcription start site +/- 1 kb). Only associations with FDR < 0.05 and at least 5 overlapping loci were considered for further analysis. To determine potential evidence in the literature for these associations, we annotated reported cases where the virus encoding for the vTR was associated with the corresponding disease (**Data S7**).

To assess whether hTFs or cofactors (CoFs) may mediate the influence of vTRs on disease-associated loci, we applied RELI to publicly available hTF/CoF ChIP-seq peak sets performed in HEK293 (or related) cells. For each hTF/CoF, RELI was used to quantify the enrichment of GWAS risk loci within ChIP-seq peaks. As in the vTR–DEG analysis, risk loci were expanded to include variants in linkage disequilibrium with published GWAS variants, and enrichment was evaluated by comparing the observed overlap to a null distribution generated through permutation using matched control regions. Significant associations were defined as hTF/CoF–disease pairs with p < 0.01. Next, we integrated these results with the vTR-disease associations identified in the DEG-based RELI analysis. For a given disease, we identified hTFs or CoFs that showed significant enrichment at the same GWAS loci for which a vTR was also significantly associated. To determine whether the vTR could potentially modulate these loci through recruitment by the hTF/CoF, we required evidence of a PPI between the vTR and the corresponding hTF/CoF. We annotated all resulting trios where (i) the vTR DEGs were significantly associated with the disease, (ii) the hTF/CoF ChIP-seq peaks overlapped GWAS loci for that disease, and (iii) a PPI existed between the vTR and the hTF/CoF (**Data S8**).

#### Comparison between hTF and vTR M1H activity

The transcriptional activities of vTRs were compared to those of hTFs tested by Lambourne et al. using the low-background M1H system.^75^ The distribution of log2(fold change) for all vTRs tested using the minimal promoter construct described previously was compared to those of reference hTF isoforms previously tested using the same reporter construct. Distributions were compared in Prism (Graphpad) using the two-tailed Mann Whitney U-test.

#### M1H activity of vTRs with different expression timing

Annotation of the expression timing of vTRs was performed via literature curation (**Data S9**). Taking into account the varied expression kinetics of different viruses, we broadly defined expression timing as follows: Immediate Early (IE) – expressed before replication of the viral genome. Early (E) – expressed following immediate-early and before replication of the viral genome. Late (L) – expressed following replication of the viral genome. Early-Late (E/L) expressed both prior to and following viral genome replication. Latent (Lat) – expressed primarily or exclusively during viral latency. The transcriptional activities of vTRs expressed at different timepoints of infection were compared using Prism (Graphpad) via Brown-Forsythe and Welch multiple ANOVA.

#### PROD-ATAC Screen

PROD-ATAC data analysis (of both the single-cell ATAC libraries and genotyping dial-out libraries) was performed on the University of Wisconsin-Madison’s Center for High Throughput Computing cluster as previously described accessed May 2025.^127^ Briefly, single-cell sequencing datafiles were converted to FASTQ format using cellranger-atac mkfastq (10x Genomics, version 2.1.0) and aligned to the hg38 reference genome via cellranger-atac count. Fragment files generated via cellranger-atac count (along with associated cell-assignment file outputs) were used to build Arrow files in ArchR. ArchR (version 1.0.3) was used for calling vTR marker peaks and performing downstream analysis using the same filter settings and quality cutoffs as described in depth previously.^127^ Briefly, all nuclei used for PROD-ATAC data analysis were first filtered for those with TSS enrichment scores ≥7 and ≥40,000 fragments. We then generated pseudobulk samples representing each vTR genotype and created in silico replicates to perform statistical tests to identify differentially accessible peaks. The addGroupCoverages command (minCells□=□20, maxCells□=□500, minReplicates□=□2, maxReplicates□=□5) merged cells of a known genotype. The resultant coverage file was queried to identify peaks via addReproduciblePeakSet. MACS3 was used with default parameters for peak calling. Patterns of motif enrichment within these differentially accessible peaks were identified by calling peakAnnoEnrichment with default parameters. Scripts relevant for downstream PROD-ATAC data processing and figure generation used in both this and other PROD-ATAC associated publications are deposited in GitHub (https://github.com/mfrenkel16/OncofusionPRODATAC/).

#### Comparing hTF/CoF interactions between viral proteins and vTRs

Protein interaction partner types for vTRs and non-vTR viral proteins were compared by taking the union of our Y2H results and interactions compiled by the Human-Virus Interaction DataBase (HVIDB) (**Data S10**).^22^ Protein interaction partners of vTRs were categorized into eight categories: hTF, cofactor, RNA processing, proteostasis, signaling, metabolism, immunity, or other (**Data S10**). The TF list was obtained from Lambert et al.,^79^ whereas cofactors include proteins that were not classified as hTF but are annotated in the list of human cofactors from Animal TF DB.^137^ RNA processing, proteostasis, signaling, metabolism, and immunity were non-hTF or cofactor proteins that were defined based on gene ontology (GO) terms generated by UniProt and confirmed via PubMed literature search. Example GO terms for each category are as follows: RNA processing - regulation of RNA splicing (GO:0043484), mRNA processing (GO:0006397), mRNA stabilization (GO:0048255), regulation of transcription by RNA polymerase II (GO:0006357). Proteostasis - protein ubiquitination (GO:0016567), protein folding (GO:0006457), protein stabilization (GO:0050821), intracellular protein transport (GO:0006886),. Signaling - signaling (GO:0023052) and related lower terms as described in Lambourne et al.^75^ Metabolism - fatty acid biosynthetic process (GO:0006633), glycolytic process (GO:0006096), cellular respiration (GO:0045333), ATP biosynthetic process (GO:0006754), TOR signaling (GO:0031929). Immune - innate immune response (GO:0045087), adaptive immune response (GO:0002250), inflammatory response (GO:000695), defense response to virus (GO:0051607). Proportions of interactions with hTF, cofactor, or other (all non-hTF or cofactor proteins) were then compared between vTRs and non-vTR viral proteins using a permutation test of a χ² statistic; two-sided empirical P values were computed.

#### Enrichment analysis for different PPI partners across vTR timing

To assess how different categories of protein interaction partners vary across vTR timing groups, we first normalized raw interaction counts to per-condition proportions. Proteins were annotated into functional categories (e.g., hTFs, cofactors, RNA processing, proteostasis, signaling, metabolism, immunity, and other). For each group, we aggregated proportions across proteins and compared them to the background average across all conditions. Fold-enrichment values were calculated, log□-transformed, and visualized as heatmaps with a fixed scale (±1.5). This approach highlights partner categories enriched or depleted in specific vTR timing groups relative to the overall interaction distribution.

#### Comparison between vTR and hTF PPI partners

Similarity of PPI partners between vTRs and hTFs was assessed by comparing the union of hTF–human protein interactions supported by at least one piece of evidence in BIOGRID^138^ or reported in HuRI^128^ with the union of vTR–human protein interactions detected by Y2H (as described) or reported in HVIDB.^22^ For each protein partner represented, we calculated the percentage of hTFs or vTRs with which it interacted. Proteins interacting with more than 5% of hTFs or vTRs were highlighted. Jaccard indices were calculated to quantify the overlap of PPI partners: within vTRs, within hTFs, and between vTRs and hTFs.

#### Determining structural similarity of homolog vTRs

Structural similarity of homolog vTRs was determined using NCBI blast to compare amino acid identities on a pairwise basis. Amino acid sequences were chosen based on the representative UniProt ID for that given vTR. Amino acid % identity values were rounded to the nearest tenth place.

## Notes

### Competing Interest Statement

The authors have declared no competing interest.

